# Contrasting behavioral and physiological effects of *Gtf2i* duplication and deletion in mouse models of the 7q11.23 Duplication and Williams-Beuren Syndromes

**DOI:** 10.1101/2025.11.13.688274

**Authors:** David Peles, Shai Netser, Natali Ray, Taghreed Suliman, Shlomo Wagner

## Abstract

7q11.23 Microduplication Syndrome (7Dup) and Williams-Beuren Syndrome (WBS) are two ASD-related syndromes characterized by both common and contrasting symptoms, caused by either duplication or deletion of a 1.5-1.8 Mb segment in section q11.23 of Human chromosome 7, respectively. Notably, WBS patients show reduced social fear and are considered hyper-social, while 7Dup patients suffer from social anxiety and withdrawal. Previous work suggests that the *GTF2I* gene, one of the genes included in this segment, has a major role in the social symptoms of both syndromes. Here, we combine video and thermal imaging in multiple social behavior tests to screen for behavioral and physiological variables showing variations in mice models with either a duplication (*Gtf2i*^+/dup^) or a deletion (*Gtf2i*^+/del^) of the gene. Our analyses of social behavior, micturition, and defecation patterns identify several differences between wild-type and mutant littermates, some of which show contrasting variations associated with *Gtf2i* dosage. Interestingly, thermal imaging revealed that *Gtf2i* dosage dictates the mice’s surface temperature profile during the tests, with *Gtf2i*^+/dup^ males exhibiting higher surface temperature than their wild-type littermates, while *Gtf2i*^+/del^ males and females show the opposite tendency. These results suggest that the two mouse models exhibit opposite changes in either their emotional state or thermoregulation capabilities, in correlation with *Gtf2i* dosage.

## BACKGROUND

7q11.23 Duplication Syndrome (7Dup) and Williams-Beuren Syndrome (WBS) are two rare genetic conditions caused by either duplication or deletion, respectively, of typically 1.5-1.8 Mb in section q11.23 of chromosome 7 (1–3). These syndromes are characterized by both common and contrasting symptoms. Most notably, social anxiety and withdrawal are common in 7Dup patients (1), while WBS patients tend to have an overly social personality and no social anxiety (3) and are generally considered as hyper-social (4). This suggests a dose-dependent effect on social behavior, caused by one or more of the involved genes. Of the various (typically 25-27) genes included in the 7q11.23 genomic section, *GTF2I* was suggested to have a pivotal role in dictating behavioral and cognitive phenotypes in both 7Dup and WBS patients, based on rare human atypical small deletion or duplication cases that include the GTF2I gene (5–8). This gene encodes the signal-induced multifunctional transcription factor II-I (TFII-I), known to be involved in a variety of gene regulatory processes (9).

Mouse models with either three copies or a single copy of *Gtf2i* have been previously used as animal models of the 7Dup and WBS syndromes (9–13). Multiple studies examined the behavioral phenotypes of such models, using mainly the three-chamber test (14). Most studies exploring the phenotype of *Gtf2i^+/Del^* mice (Del mice, for short) reported a hyper-social behavior (13, 15, 16). In contrast, studies of *Gtf2i^+/Dup^* mice (Dup mice, for short) reported less consistent results, with one study reporting deficits in social behavior, including loss of sociability and social novelty preference (12), and another study reporting an increased number of ultrasonic vocalizations (USVs) made by pups following separation from their mother (10). Other studies, however, did not find deficits in the social behavior of Dup mice (16). It should be noted that the only studies directly comparing the behavioral phenotype of both Dup and Del mice found no statistically significant contrasting effects of the opposite types of *Gtf2i* copy-number variation (CNV) (10, 16).

Here, we combined a battery of social behavior tests with a multi-modal analysis to directly compare the phenotypes of Dup and Del mice and their WT littermates. Besides stimulus investigation behavior, these modalities include micturition and defecation patterns and changes in surface temperature at various body parts, all measured using infrared thermography (IRT), a non-invasive method previously used in mice to assess anxiety and stress (17–19). Specifically, we looked for behavioral and physiological variables showing contrasting changes between the two models, thus suggesting dependency on *Gtf2i* dosage. After normalizing the results of each type of mutant mice (Dup and Del) with their WT littermates (WT_Dup_ and WT_Del_, respectively), we found that Dup male mice exhibit reduced social discrimination capabilities as compared to Del male mice. For micturition and defecation behaviors, we found a higher rate in Dup male mice than in Del male mice, mainly during the Social Preference (SP) test. Finally, Dup and Del mice showed contrasting differences in surface temperature in comparison to their WT littermates, with Dup male mice showing higher surface temperature of their eyes, body center, and at the base of their tail, while Del male mice showing lower surface temperature at their body center and at the base of their tail. These results suggest opposite changes in either the emotional state of the two types of mutant mice, or their thermoregulation, in correlation with *Gtf2i* dosage. Altogether, our study identifies several specific behavioral and physiological variables, including surface temperature, which vary in correlation with *Gtf2i* dosage, hence are likely to be dictated by the *Gtf2i* gene.

## MATERIALS AND METHODS

### Animals

Subject mice were wild type (WT) or mutant adults (12-20 weeks old) with a CD1 (ICR) genetic background, derived from breeding either a *Gtf2i*^+/dup^ (Dup) or a *Gtf2i*^+/Del^ (Del) male mouse (10) with a CD1 female. Subject mice were bred and raised in the SPF mouse facility of the University of Haifa. Stimuli mice were either juvenile (3-6 weeks old, used for the social preference (SP) test, see below) or adult (for the Sex Preference (SxP) and Emotional State Preference (ESP) tests) CD1 mice. CD1 mice were purchased from Envigo (Rehovot, Israel) and held afterwards in an SPF mouse facility at the University of Haifa. Mice were housed in groups of 3-5 in a dark/light cycle of 12 hours (Lights on at 9 pm) with *ad libitum* food and water and veterinary supervision. Experiments were conducted in the dark phase of the dark/light cycle. All experiments were approved by the University of Haifa Institutional Animal Care and Use Committee (IACUC) (Reference #: UoH-IL2301-103-4).

### Setup and Video Acquisition

Our setup is based on the setup previously described in (20) and (21). Briefly, a black plexiglass box arena (37cm x 22 cm x 35 cm) was placed in a sound-attenuated chamber. A visible-light (VIS) camera (either Flea3 or Grasshopper3 USB3.0, both of Teledyne FLIR) with a wide-angle lens and 30 frames per second video acquisition rate, and an infrared (IR) thermal camera (Thermapp MD, Opgal) with 8.66 frames per second were placed about 70 cm above the arena’s floor (Fig. 1A). The IR camera outputs apparent temperature per pixel, which was captured using the manufacturer’s SDK (version: EyeR-op-SDK-x86-64-2.15.915.8688-MD) and previously published Python scripts (21). This camera was designed to measure human skin temperature and assumes human skin emissivity. To improve image quality and pixel uniformity, the IR camera was turned on at least 15 minutes before starting any experiment. A high-emissivity black body (Nightingale BTR-03, Santa Barbara Infrared Inc.) was set to 37°C and was placed in the field of view of the IR camera, in order to compute a correction offset for the apparent temperature measurements, as previously described (21).

**Fig. 1.**
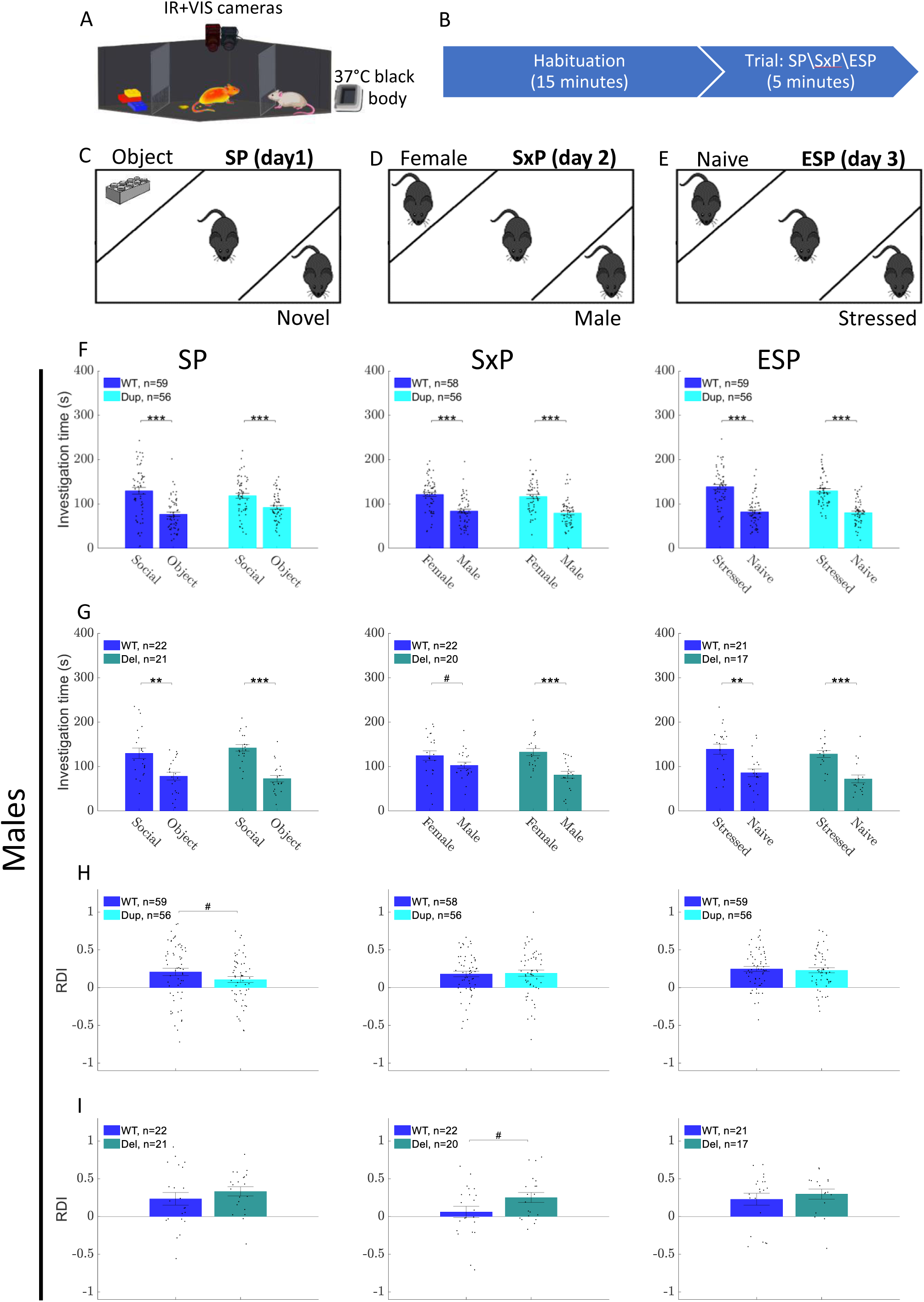
Gtf2i dosage affects the level of social discrimination in male mice. (**A**) The experimental setup includes an arena with two stimuli at opposite corners. The subject mouse is imaged by a visible wavelength camera and a thermal camera, which are positioned above the arena. A high emissivity black body set to 37°C is used as a reference to improve apparent temperature measurement from the thermal imaging. (**B**) The experiments include a 15-minute habituation period followed by a 5-minute trial period in which stimuli are positioned at the corners of the arena. In Social Preference (SP) (**C**), a toy is positioned in a chamber on one side of the arena while a novel sex-matched stimulus mouse is positioned at the other side. In the Sex Preference (SxP) (**D**), the stimuli are male and female mice. In the Emotional State Preference (ESP) (**E**), the stimuli are a naïve mouse and a stressed mouse. (**F, G**) Shows the mean investigation time (±SEM) towards each of the two stimuli during SP, SxP, and ESP of WTDup vs Dup (**F**), and WTDel vs Del (**G**). (**H, I**) Show the mean RDI (±SEM) towards stimulus1 (Social in SP, Female in SxP, and Stressed in ESP) for WTDup vs Dup (**H**), and WTDel vs Del (**I**). A two-sided Wilcoxon rank-sum test was used for pairwise comparisons. (**F,G**) were FDR corrected using the Benjamini-Hochberg method (2 comparisons per panel). p-values equal to or smaller than 0.1,0.05,0.01, and 0.001 were marked by #, *, **, ***, respectively.

### Social discrimination tests

We used three social discrimination tests, as previously described in (21) and (22), each performed on a different day. All tests included a 15-minute habituation stage in which the mouse got used to the arena. During the habituation period, two empty triangular chambers, each with a metal mesh (18cm width x 6cm height with 1x1 cm holes) at its bottom, allowing the subject mouse to investigate its content, were located in two opposite corners of the arena. The habituation stage was followed by a 5-minute trial stage in which the empty chambers were replaced with stimulus-containing chambers: a sex-matched juvenile mouse vs. a Lego toy in the SP test, an adult male vs. an adult female in the SxP test, and a sex-matched adult mouse stressed by being located for 15 minutes within a restrainer before the test vs. a sex-matched adult naïve mouse in the ESP test. All social stimuli were novel to the subject mouse (Fig. 1 B-E). To assess the effect of stress on the subject mice’s temperature, we used a fourth test, which was done on a separate day (after the mice had completed the SP, SxP, and ESP tests), in which the subject mouse was held in a restrainer for 15 minutes before performing an additional SP test (Fig. 3F).

**Fig. 2.**
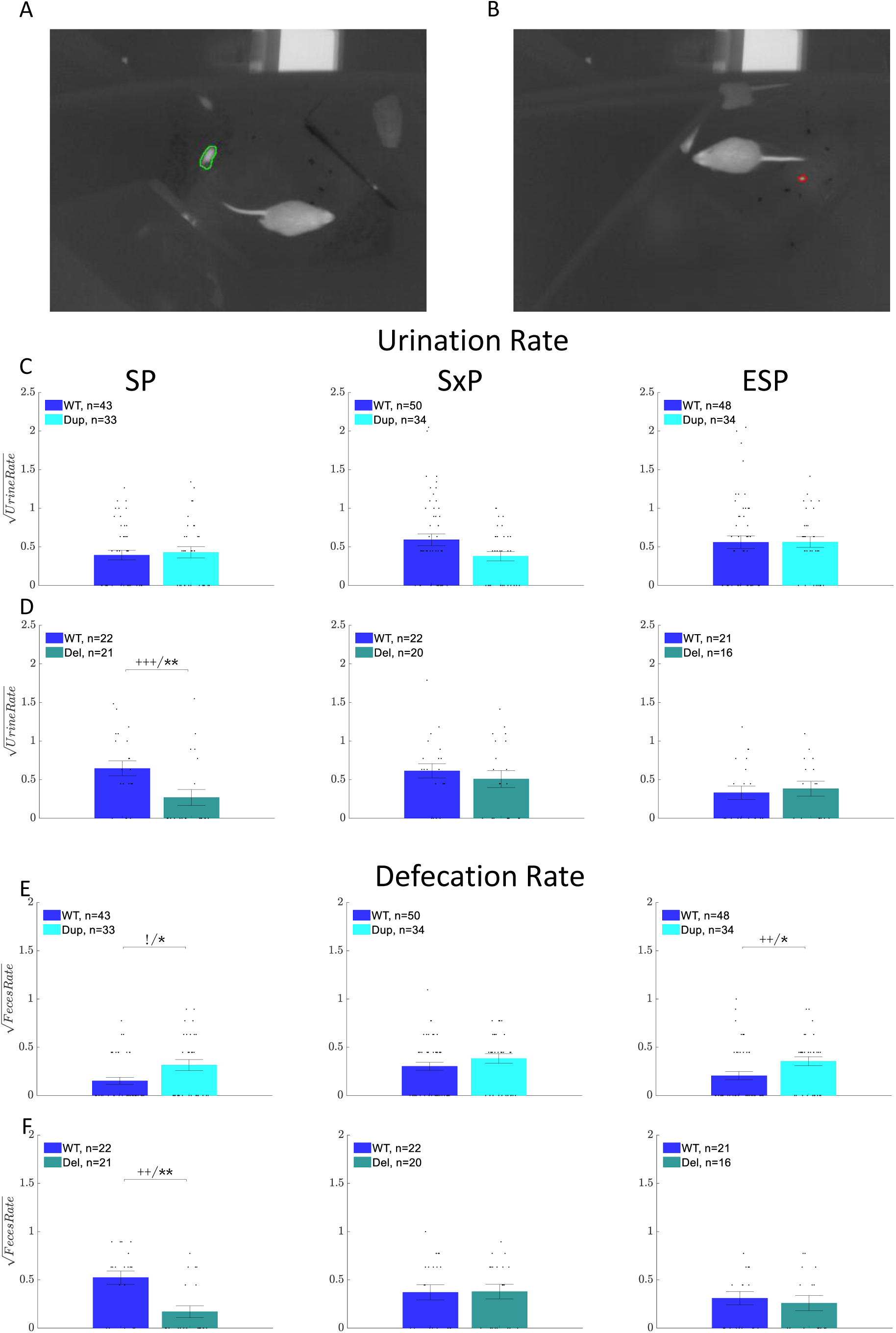
Gtf2i dosage affects urination and defecation rates in male mice. Example of an automatic detection of a urination spot (**A**) and a defecation spot (**B**) in a thermal image during a social behavior test. The detected urine segment is marked in green (**A**), and the detected feces segment is marked in red (**B**). (**C,D**) Square root of the # of urination spots per minute during SP, SxP, and ESP for WTDup vs Dup (**C**), WTDel vs Del (**D**). (**E,F**) shows the same for defecation rate. Bars show the mean value, and error bars show ±SEM. For pairwise comparisons, we used two types of tests. A two-sided Wilcoxon rank-sum test with p-values equal to or smaller than 0.1,0.05,0.01, and 0.001 was marked by #, *, **, ***, respectively. A two-way Chi-square test (for comparing the number of zeros and non-zeros between two groups) with p-value smaller than 0.1,0.05,0.01, and 0.001 was marked by !,+,++,+++, respectively.

**Fig 3.**
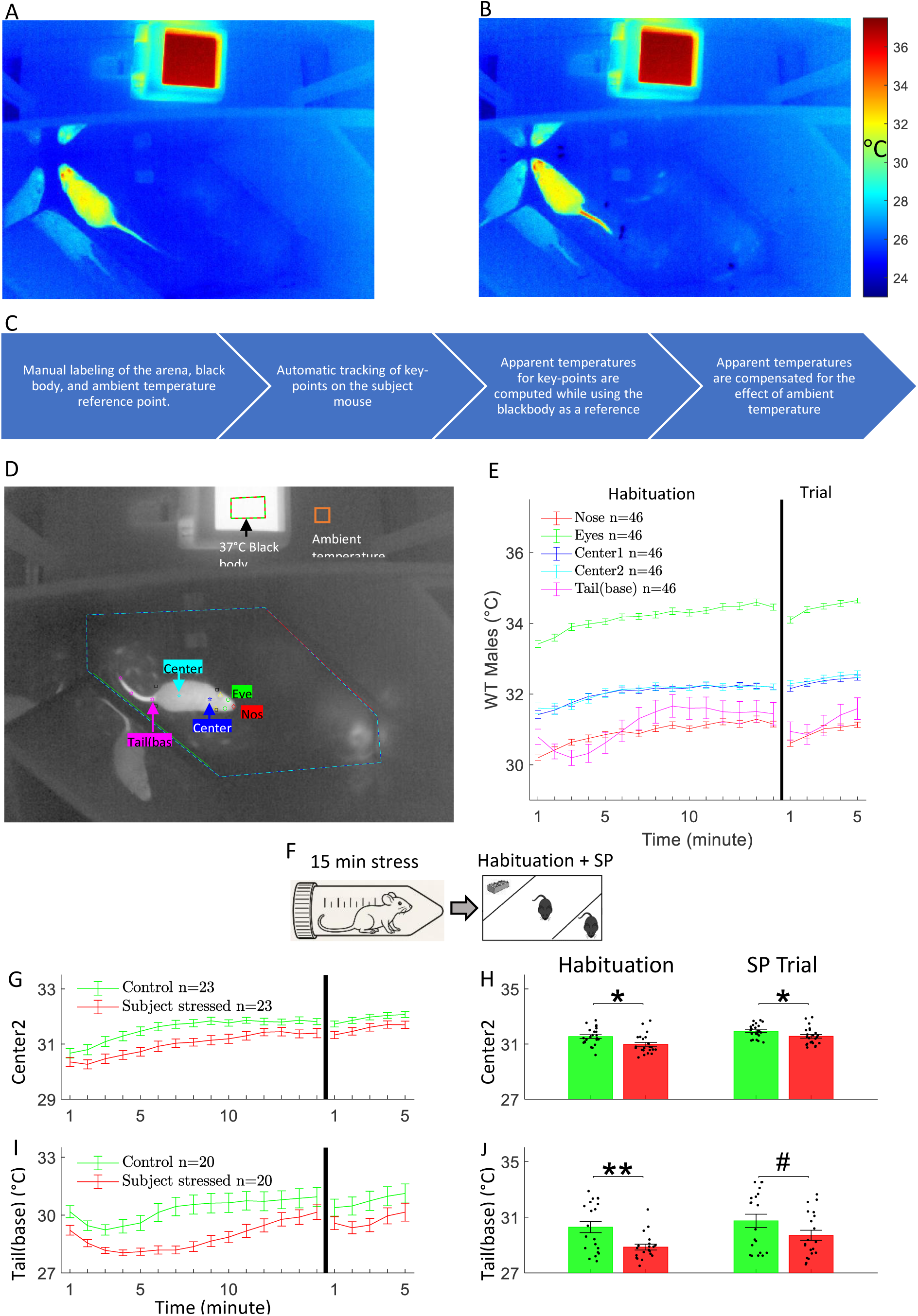
Body surface temperature changes along the time course of the experiment in a body part-dependent manner. (**A,B**) Apparent temperature images taken at 3 minutes (**A**) and 12.5 minutes (**B**) after the beginning of habituation of the same SP experiment of a WTDel male subject mouse. Differences in apparent temperature are seen in the tail and also in the back. (**C**) Workflow of the temperature analysis method. (**D**) A thermal image showing the mouse’s body parts detection by the DLC software, the manual labeling of the black body 37°C temperature reference (red-green line), the location of the ambient temperature reference point (orange rectangle), the annotation of the arena’s floor (blue-cyan line) and the annotation of the side of each stimulus, marked by a green line for stimulus1 and a red line for stimulus2. (**E**) The mean ±SEM surface temperature of several body parts across the SP test for WTDup and WTDel mice, pooled together. (**F**) The subject stress paradigm includes 15 minutes in which the subject mouse is kept in a restrainer, followed by 15 minutes habituation and 5 minutes SP trial. Stressed WTDup and WTDel animals (pooled together) exhibit lower temperature in both Center2 (**G**,**H**) and Tail(base) (**I**,**J**). A two-sided Wilcoxon rank-sum test was used for pairwise comparisons. (**H**,**J**) were FDR corrected (2 comparisons per panel). p-values equal to or smaller than 0.1,0.05,0.01, and 0.001 were marked by #, *, **, ***, respectively

### Analysis of investigation behavior

Analysis of the VIS videos was done using TrackRodent software as previously described (20, 23). For each subject mouse, we measured the total time spent by the subject on investigating each stimulus and computed the Relative Discrimination Index (RDI), defined as the difference in investigation time between the two stimuli, divided by their sum. A positive RDI value shows a preference for stimulus 1, which is considered to be the social stimulus in the SP test, the female in the SxP test, and the stressed mouse in the ESP test, while a negative RDI value shows the opposite preference.

### Analysis of the thermal videos

A previously published graphical user interface (GUI) (21) was used to annotate the thermal videos. Specifically, it was used to annotate the arena’s floor, mark the side of each stimulus, mark the location of the black body surface, and specify the first and last frames of the habituation period and the trial period of each session. In addition, a location on the plastic shelf that holds the arena was marked and used to compensate for the ambient temperature (Fig. 3D).

### Analysis of urination and defecation activities

Urination and defecation rate per minute were computed using DeePosit software, as previously described (21). Videos with a significant occlusion of any part of the arena were excluded (55 of 767 videos were excluded, see Supplementary Table 1). We considered an occlusion to be significant if part of the mouse’s body (and not just its head) was occluded in parts of the video.

### Analysis of body surface temperature

#### Tracking the subject’s body parts in thermal videos

Key-points marking the body parts of the subject mouse were detected in each IR image using the DeepLabCut (DLC) software (24) version 3.0.0rc4. In order to feed the videos to the DLC software, the recorded thermal videos were converted to 8-bit per pixel AVI video files by linearly mapping the recorded apparent temperature range [15°C .. 40.5°C] to the grayscale range [0 .. 255], resulting in 0.1°C thermal resolution. For training the DLC algorithm, 1999 IR images were manually labeled using DLC’s GUI. Labeled body parts included: nose, left eye, right eye, left ear, right ear, center1 (the neck of the mouse), center2 (the body center), tail-base, tail middle, tail end, and four limbs (See Fig. 3D). Of all these key points, we analyzed only the nose, eyes (max of both), center1, center2, and tail-base. The DLC model training was done using default parameters of the PyTorch-based engine for 200 epochs, with a batch size of 32 and ResNet50 backbone (25).

After training, DLC was activated on all IR frames of habituation and trial across all valid experiments. To prevent DLC from detecting key points on the body of the stimulus mouse (which was visible during the trial in some of the experiments), the area in the image that contains the stimulus (determined as the area 14 pixels away from the lines that mark the stimulus chamber’s border) was set to a constant value (matching 22°C).

#### Extracting the temperature for each body part during the experiment

The thermal camera supplies the apparent temperature for each pixel in the video, which is further corrected by subtracting the black body temperature reference and adding 37°C. To improve the reliability of the thermal measurements of the tracked body parts, frames in which the DLC grade for one of the more reliably-tracked body parts (center1, center2, or tail-base) was lower than 0.7 were ignored (DLC grade for each key point ranges between 0 and 1). In general, a temperature measurement of a specific body part in any given frame was ignored if the DLC grade of the detection of this body part was smaller than 0.8. Moreover, a body part temperature was considered valid only if it was inside the annotated arena’s floor area and at least 6 pixels away from its borders (to account for the smoothing radius, see below). Specifically, we measured the eyes and nose temperature in a given frame only if the detection of both eyes and nose at this frame was valid (was inside the arena’s floor area and 6 pixels away from its borders) and their detection grades were higher than 0.8. The tail-base temperature was determined as the maximal temperature in a radius of 6 pixels surrounding the coordinates computed by DLC (smoothing radius). This compensates for possible minor mistakes in the detected location of the tail-base, which might result in a significant temperature error due to its narrow size. For center1 and center2, we used the value of a smoothed thermal image (smoothed by a Gaussian of 11x11 pixels and std=2.5 pixels), instead of the original value, in order to reduce noise. We required each body part temperature to have at least 10 valid measurements per minute during both the habituation and trial stages in order to be included in the analysis. Videos in which the mouse had a dark spot on its fur (due to urine or feces on its fur) or fur damage were excluded (33 out of 767 videos were excluded, see Supplementary Table 1).

We found that the measured body parts’ apparent temperature was correlated to the ambient temperature in the room, which is consistent with previous literature (26, 27). We determined the ambient temperature as the mean apparent temperature of 11x11 pixels surrounding a fixed manually marked point on the plastic shelf on which the arena is positioned (see Fig. 3D). For each body part, we computed a linear regression (see Supplementary Fig. 5A) between the ambient temperature and the average body part temperature during minutes 10-14 of the habituation (when the temperature was found to be most stable, see Supplementary Fig. 5D-I). A compensation function was then used to compensate for the ambient temperature deviation from the mean ambient temperature (23.74°C) (see Supplementary Fig. 5B). The compensation function parameters were computed from the videos of all of the experiments pooled together, including males and females of all genotypes. For each body part, the relevant compensation function was used across all genotypes and experiments. Following all these compensation steps, the median value of the valid measurements of a given body part over a minute was used as the final temperature for further analysis. To compute the temperature for longer periods, we averaged the values of the relevant minutes. Throughout this work, we assumed that the mice’s fur properties are similar across subject mice, and hence, the differences in temperature measurement by the thermal camera may be attributed to the change in temperature and not to differences in the fur emissivity or density.

### Analysis of average speed per minute

We relied on the detection of Center2 by DLC in the thermal videos to compute the average speed (cm per second) of the subject mouse averaged for each minute of the habituation and trial stages. Center2 detections that had a DLC grade lower than 0.8 were ignored. The speed per second was calculated using the Euclidean distance divided by the delta time for all pairs of frames with roughly 1 second between them. Minutes with fewer than 50 valid speed measurements were ignored.

### Normalization to WT values

In many cases, we found that the WT animals of the two lines showed significant differences between them. Therefore, for a direct comparison between Dup and Del mice, we normalized the results of mutant mice to the results of their WT littermates. In most cases, this was done by dividing the results of each mutant subject animal by the mean value of the sex-matched relevant WT mice in the relevant tests and periods. When comparing normalized RDI values, however, the mean RDI values of the relevant WT mice were subtracted from the RDI values of the mutant mice.

### Statistical analysis

Pairwise comparisons were computed using a non-parametric two-sided Wilcoxon rank-sum test (using MATLAB’s *ranksum* function), with p-values equal to or smaller than 0.1,0.05,0.01, and 0.001 were marked by #, *, **, ***, respectively. In addition, for the comparison of micturition and defecation rates, we used a two-way Chi-square test to compare the number of zeros and non-zeros between the relevant genotypes, as previously described in (21). A significant difference between the number of zeros across two genotypes with p-value smaller than 0.1,0.05,0.01, and 0.001 was marked by !,+,++,+++, respectively. All panels with multiple comparisons were FDR corrected using the Benjamini-Hochberg method (28). The p-values (before and after FDR correction), additional statistical data, and the data points for all of the statistical comparisons in this work are provided in Supplementary file 1.

## RESULTS

### Social discrimination behavior negatively correlates with *Gtf2i* dosage

For behavioral phenotyping, each subject mouse performed three different social discrimination tasks in an experimental arena equipped with a visible wavelength (VIS) camera and an infra-red (IR) camera, as previously described (21, 22) (Fig. 1A). Each task comprised a 15-minute habituation stage with two empty chambers located at opposite corners of the experimental arena. After the habituation period, the two empty chambers are replaced by a similar, stimulus containing chambers for a 5-min trial period (Fig. 1B). In the SP test, conducted on the 1^st^ day of experiments, the chambers contained a sex-matched mouse vs. a Lego toy (Fig. 1C). In the SxP test, conducted on the 2^nd^ day of experiments, the chambers contained a female vs. a male mouse (Fig. 1D). In the ESP test, conducted on the 3^rd^ day of experiments, the chambers contained a stressed vs. a naïve mouse (Fig. 1E).

The time spent by the subject mouse for investigating each of the stimuli was analyzed using the VIS video clips using the TrackRodent software (20). We found that for male mice, all four genotypes (Dup, WT_Dup_, Del, WT_Del_) exhibited a clear preference to investigate the social stimulus over the object stimulus in the SP test, the female stimulus over the male stimulus in the SxP, and the stressed animal over naïve animal in the ESP test (Fig. 1F-G). Similar results were found for female mice, although in that case, WT_Dup_ did not show a sex preference at all, suggesting that WT_Dup_ and WT_Del_ animals are different in their behavior (Supplementary Fig. 1A-B). Interestingly, unlike their WT littermates, Dup female mice showed a preference towards the male mouse in SxP (Supplementary Fig. 1A). Overall, both Dup and Del mice did not show any deficit in their social discrimination behavior, compared to their WT littermates. Nevertheless, when a relative discrimination index (RDI) reflecting the preference towards stimulus 1 (social stimulus in SP, female in SxP, and stressed stimulus in ESP) was calculated for each animal, we found a borderline significant difference between Dup and WT_Dup_ male animals in the SP test and between Del and WT_Del_ male mice in the SxP test (Fig. 1H-I). Moreover, when the results of the mutant mice were normalized to those of their WT littermates, we found that the normalized time spent by Del mice on investigation of stimulus 1 in the SP test was significantly higher than in Dup mice, while an opposite difference was found for stimulus 2 (Supplementary Fig. 2A). Similar tendency, although not statistically significant was observed for SxP, while in the ESP test the difference in stimulus 2 investigation time reached borderline statistical significance (Supplementary Fig. 2A). Accordingly, the normalized RDI was significantly higher in Del than in Dup mice across SP, SxP while a non-significant trend appears also in ESP (Supplementary Fig. 2C), suggesting a higher capability of social discrimination in Del compared to Dup male mice. As for females, RDI values of Dup and Del were not significantly different from their WT littermates, except for borderline significantly lower RDI in SxP of Dup females due to higher preference to the male stimuli (Supplementary Fig. 1C-D). As for the normalized RDIs, they exhibited the same trends as males, but the only significant difference in normalized values between the mutant animals was in the SxP test, where Dup females showed higher investigation than Del females, which was also reflected by their higher normalized RDI values (Supplementary Fig. 2B, D). Taken together, these findings suggest that in male mice, *Gtf2i* dosage negatively correlates with the level of social discrimination, in accordance with the hyper/hypo-social phenotype observed in humans.

### Gtf2i dosage affects micturition and defecation patterns during social behavior

Another aspect of social behavior in mice is territorial scent marking, carried out by micturition and defecation activities (29, 30). We previously presented a computational tool, termed DeePosit, for AI-based automatic detection and analysis of micturition and defecation activities during behavioral tasks in mice from thermal video clips (21). Here, we used DeePosit to analyze these activities during the three social behavior tasks described above, across all four genotypes (Fig. 2A-B). For micturition in male mice, we found a significantly lower rate in Del compared to WT_Del_ mice during the SP task, while no significant difference was found between Dup and WT_Dup_ male mice (Fig. 2C-D). No difference was found for both Dup and Del females (Supplementary Fig. 3A-B). As for defecation activity, we observed a significantly higher rate in Dup males during the SP and ESP tasks and an opposite change in Del mice specifically during the SP task, compared to their WT littermates (Fig. 2E-F). Accordingly, after normalization to WT values, Dup mice showed significantly higher rates of both micturition and defecation, compared to Del mice, in the SP task and of defecation in the ESP task (Supplementary Fig. 4A,C). No difference between mutant and WT mice was found for females in urination and defecation activity (Supplementary Fig. 3, Supplementary Fig. 4B,D), suggesting that the differences between the lines in both micturition and defecation are sex-specific. Assuming that micturition and defecation activities are associated with territorial behavior, these results may suggest a higher territoriality during social interactions in male Dup mice, which fits the social withdrawal symptoms characterizing the 7Dup syndrome.

### *Gtf2i* copy number variation affects mouse surface temperature in a dosage-dependent manner

Thermography is a non-invasive method shown to be efficient for assessing affective states in various animal species (31). In this method, a thermal camera is used for assessing the surface temperature of the body, which is influenced by two contrasting processes: (1) a rapid arousal-induced vasoconstriction that cools the animal’s skin, and (2) a slow stress-induced hyperthermia that heats it (31, 32). We therefore analyzed the surface temperature in several body parts of subject mice from the thermal video clips taken by the IR camera during the experiments described above (Fig. 3A-B). For that, we combined DeepLabCut (DLC), a machine learning based key point detector (24), with the processing pipeline shown in Figure 3C. Shortly, following manual marking of the arena’s floor, the 37°C black body and a spot on the plastic shelf that holds the arena (termed “ambient temperature reference point”), we employed DLC to extract the location of the following body parts: Nose, Eyes, Center1 (the back of the neck), Center2 (the body center) and Tail(base) (Fig. 3D). The apparent temperature for each body part was compensated for the ambient temperature using a linear regression (Supplementary Fig. 5A-C), see methods for more details.

We found significant changes in the apparent temperature of the various body parts along the time course of the experiments, with the most prominent changes observed in the tail(base) (Fig. 3E and Supplementary Fig. 5D-I). Accordingly, we found a distinct profile of temperature changes for each body part, with slight differences between males and females (Supplementary Fig. 5D-I). Generally, most body parts showed a steady increase in temperature along the first 5-10 minutes of habituation, which might be related to stress-induced hyperthermia caused by the introduction of the subject to the arena. In contrast, the Tail(base) exhibited a unique pattern of surface temperature changes, comprised of an initial decrease at the first three minutes of habituation, followed by an increase during the next five minutes and stability through the rest of the habituation stage. Following the introduction of stimuli at the beginning of the trial stage, most regions exhibited a transient decrease in their apparent temperature, most probably due to a transient arousal induced by stimuli introduction. The profiles of the various body parts were apparently preserved between the various tasks (Supplementary Fig. 5D-I), with the exception of the Tail(base), which got gradually hotter from day 1 (SP) to day 3 (ESP), suggesting that it might be especially sensitive to the arousal state induced by the transfer to the arena. This arousal state is expected to gradually decrease between the days of the experiment, as the animal gets familiar with the arena and procedure.

To assess the influence of affective states on the measured surface temperature, we repeated the SP test on a separate day after stressing the subject mouse by confinement in a restrainer for 15 minutes before the test (Fig. 3F). Stressed mice showed reduced surface temperature in both Center2 and Tail(base) regions and to a lower extent in Center1, but not in the eyes and nose except for some trend in the first minute of the habituation (Fig. 3G-J and Supplementary Fig. 6). Thus, Center2 and Tail(base) areas seem to be most sensitive to the affective state of the mouse.

For comparing the surface temperature between the various genotypes, we focused our analysis on Center2, Tail(base), and Eyes. When pooling together the results across all three tests (see Supplementary Fig. 7-10 for the results of each test separately), we found that Dup male mice display a higher temperature, compared to their WT littermates, at all three regions, throughout the habituation and trial stages in a statistically significant manner (except for Center2 during the trial stage where we observed a non-significant trend; Fig. 4A-F). In contrast, Del male mice had a lower temperature than their WT littermates throughout the habituation and trial stages in both Center2 and Tail(base), while no significant difference was found in Eyes. (Fig. 4G-L). When comparing Dup and Del male mice after normalization with their WT littermates, we found that Dup mice are significantly warmer than Del mice in all tests and across all test stages except for the Eyes during the ESP trial, where only a non-significant trend is visible (Supplementary Fig. 7E, 8E, 9E). Interestingly, a different pattern of differences between mutant and WT mice was found for females, with both Dup and Del females showing lower Tail(base) temperature and no difference in Eye temperature, compared to their respective WT animals (Fig. 5). In Center2, however, only Del females exhibited lower surface temperature compared to their WT littermates, while no difference was found in Dup females (Fig. 5B,E,H,K). After normalization to the relevant WT, Dup female mice had higher Center2 temperature than Del female mice in SP and ESP, and in Tail(base) during all tests (Supplementary Fig. 7F, 8F, 9F). It should be noted that we did not find differences in the average movement speed of the mice that may explain the higher temperature in Dup mice or lower temperature in Del mice (Supplementary Fig. 11), suggesting that the thermal difference is not due to physical exertion.

**Fig. 4.**
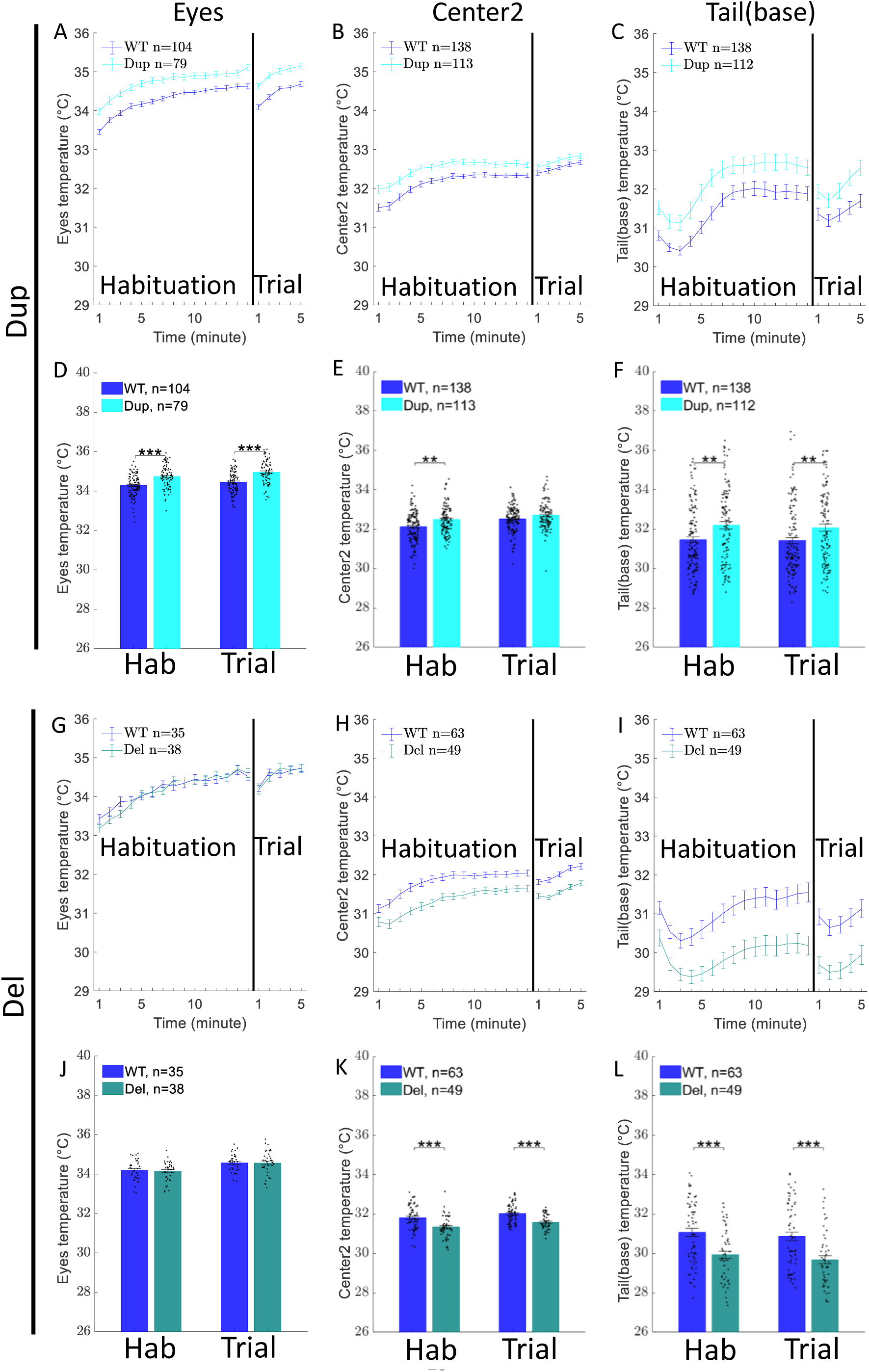
Surface temperature varies in contrasting directions in Dup and Del male mice, compared to their WT littermates. Mean ±SEM of Eyes, Center2, and Tail(base) temperature during the three behavioral tests (pooled together) for male mice in habituation minutes 1-15 and trial minutes 1-5 for Dup vs WTDup (**A-F**) and Del vs WTDel (**G-L**). Pairwise comparisons in (**D**-**F,J-L**) were done using a two-sided Wilcoxon rank sum test, which were FDR corrected (2 comparisons per panel). p-values equal to or smaller than 0.1,0.05,0.01, and 0.001 were marked by #, *, **, ***, respectively.

**Fig. 5.**
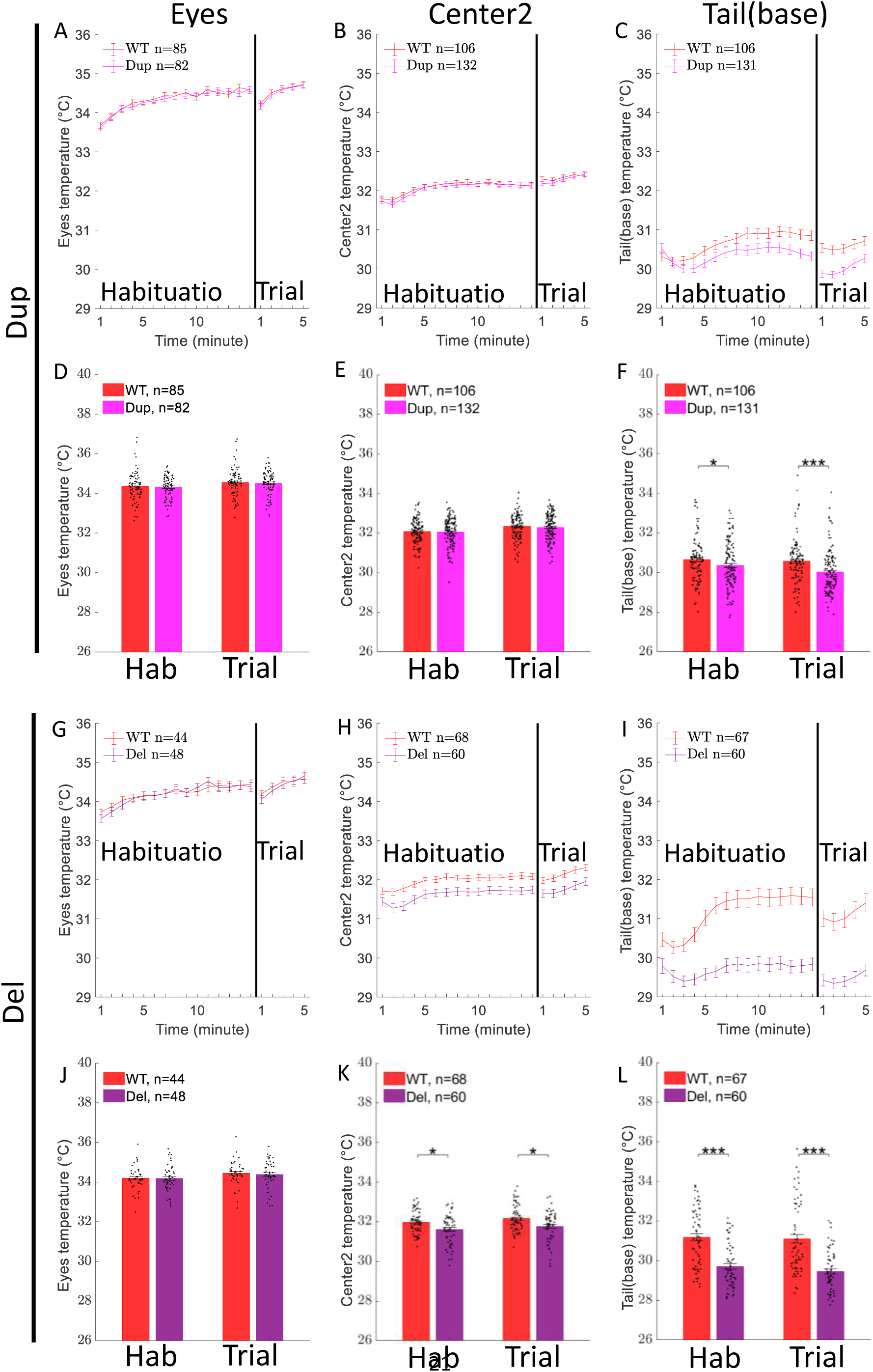
Surface temperature varies in Dup and Del female mice, compared to their WT littermates. Mean ±SEM of Eyes, Center2 and Tail(base) temperature during the three behavioral tests (pooled together) for female mice in habituation minutes 1-15 and trial minutes 1-5 for Dup vs WTDup (**A-F**) and Del vs WTDel (**G-L**). Pairwise comparisons in (**D**-**F,J-L**) were done using a two-sided Wilcoxon rank sum test, which were FDR corrected (2 comparisons per panel). p-values equal to or smaller than 0.1,0.05,0.01, and 0.001 were marked by #, *, **, ***, respectively.

Overall, we found that *Gtf2i* dosage affects the apparent surface temperature of the Dup and Del male and female mice.

## DISCUSSION

The GTF2I gene, located in the 7q11.23 region of the human genome, is thought to have a critical role in causing the opposite effects on social behavior of the Williams-Beuren syndrome (WBS) and the 7Dup syndrome (5–8, 33). Here, we conducted a multimodal phenotyping of the social behavior of Del (*Gtf2i^+/Del^*) and Dup (*Gtf2i^+/Dup^*) mice, two mouse models of the WBS and 7Dup syndromes, respectively. For that, we analyzed a variety of behavioral and physiological variables recorded while the animals performed a battery of three social discrimination tests: SP, SxP, and ESP (34). We compared the results between the mutant mice and their WT littermates and between the two mutant lines after normalization to their respective WT animals. It should be noted that WT_Dup_ and WT_Del_ mice differed significantly in some of the variables. These differences may be due to subtle variations in their genetic background, kept despite the fact that we continuously bred both lines with CD1 mice for several generations. Thus, a direct comparison with their WT littermates, as well as the normalization of the mutant mice results to the relevant WT mice results, as we did here, was essential.

We found that all genotypes showed a significant preference in all three social discrimination tests (SP, SxP, and ESP). While our experimental system and behavioral paradigm are not identical to the three-chamber test, the SP test may be considered analogous to the sociability stage of the three-chamber test (20, 35). Thus, our results are in agreement with the multiple previous studies showing proper sociability in both Del and Dup mice (13, 15, 36), but in odds with Lopez-Tabon et al (12), who reported a loss of sociability in Dup mice. Nevertheless, we found that after normalization to WT values, Del mice exhibited a higher level of preference compared to Dup mice, as reflected by their RDI values, and this difference was consistently significant across the SP and SxP tests (Supplemental Fig. 2C). Thus, in agreement with a previous study comparing the behavior of Dup and Del mice (16). In our hands, Del mice showed a better social performance than Dup mice, which is in accordance with the contrasting phenotypes of the WBS and 7Dup syndromes observed in humans (1, 3, 5, 33).

We used our thermal imaging for two purposes. First, we used it, in combination with our DeePosit tool (21), to analyze the micturition and defecation activities during the trial stage of the various tests (21). We found that Del male mice showed a lower micturition rate during the SP trial (Fig. 2D). As for defecation, we observed a significantly higher defecation rate during the SP trial in Dup mice and the opposite trend in Del mice. The higher level of both micturition and defecation rates in Dup mice may reflect stronger territorial behavior due to a higher level of social anxiety, relative to Del mice. This may explain why these differences between the genotypes were observed mainly during the SP test, which was done on the first day of the experiments. On that day, when the animals are not yet familiar with the arena and paradigm, they are expected to exhibit a higher level of anxiety compared to the next two days of experiments. It should be noted that gastrointestinal activity was shown to be affected by stress in mice (37) and that stress was shown to increase defecation rate in rats (38). It is also interesting to note that gastrointestinal issues are common in both 7Dup and WBS patients (1, 39). Together, our analyses of micturition and defecation activities suggest a higher level of social anxiety in Dup mice, as compared to Del mice, in accordance with the differences in social anxiety characterizing the WBS and 7Dup syndromes (1, 3, 5, 33).

Finally, we used thermal imaging for IRT in order to investigate the surface temperature of the mice during the various experiments. Changes in this surface temperature were shown to reflect the affective state of the animals due to the contrasting effects of the rapid arousal-induced vasoconstriction in the periphery and slow stress-induced hyperthermia. These processes lead to skin temperature changes, especially in areas with a higher density of arteriovenous anastomoses (AVAs) (see (31) for a comprehensive review). By combining DLC analysis with the thermal clips, we could analyze the dynamics of temperature changes in the various body locations along the time course of the three behavioral tests. In accordance with previous studies, we found that the tail showed different dynamics of surface temperature than the other body parts (19, 32, 40, 41). Specifically, while most parts exhibited a gradual increase in temperature along the habituation stage, the tail showed a decrease at the first three minutes and then exhibited a rapid temperature increase till reaching a stable level after about five more minutes. Similar distinct profiles of surface temperature changes following transfer of the animals to a new environment were previously reported by multiple studies in both mice and rats (17, 18, 42, 43), and seem to be related to the AVA tissue in the tail of rats and mice. Interestingly, at the beginning of the trial stage, when stimuli were introduced to the arena, we observed a transient drop in the surface temperature at most body parts, especially the tail, eyes, and nose. This seems to reflect a transient arousal state of the animal, induced by the stimuli introduction. It should be noted that a similar transient drop in surface temperature was previously reported in rats following several arousing stimuli, including the introduction of a social stimulus (44).

For comparison between the various genotypes, we chose the Eyes, body center (Center2), and the beginning of the tail – Tail(base) as each of these regions showed a distinct profile of thermal dynamics. Surprisingly, we found that *Gtf2i* affects male mice’s surface temperature during social behavior tests in a *Gtf2i* dosage-dependent manner, with Dup mice showing higher temperature than their WT littermates, especially during habituation, in all three body parts, while Del mice showing the opposite change in Center2 and Tail(base) but not in the Eyes. Notably, introducing stress to the subject mice resulted in lower Center2 and Tail(base) temperature, suggesting that lower temperature in these regions is indicative of stress. Additionally, a positive gradual increase in the Eyes temperature was seen during the habituation and trial periods in all tests, suggesting that the eyes temperature better reflects the arousal-induced hyperthermia, while the Center2 and Tail(base) are affected by both stress-induced vasoconstriction and arousal-induced hyperthermia. Notably, the contrasting trend of changes in surface temperature was consistent across all tests in males, while in females, this effect was found significant for Tail(base) in all tests and in Center2 in SP and ESP. Since we did not find differences between the genotypes in the movement speed during the various experiments, which may explain the temperature difference, the surface temperature difference might reflect variations in their arousal or stress level (17–19) or might be the result of other mechanisms, such as thermoregulation. Interestingly, differences in thermoregulation were found using rectal temperature measurement in *Shank3* mutant mice, which responded differently than WT animals to LPS injection. IRT was used in humans for assessing emotional state and mental health conditions (45). Ganesh et al. (46) used IRT to classify thermal images of ASD children and achieved an accuracy of ∼90%. They found higher facial skin temperature most prominently at the nose tip in ASD children that were presented with visual emotional stimuli. Similarly, higher nose tip temperature was found by Fernández et al. (47) in people with higher social anxiety during social behavior tasks.

### Limitations

One limitation of our study is that it does not include measurement of the basal temperature of the mice in the home cage, so we cannot say if the temperature difference we found reflects a different response to the experiment or a different basal temperature difference that is caused by the *Gtf2i* dosage. However, we did show that strong stress results in a decrease of the Center2 and Tail(base) temperature, probably due to vasoconstriction occurring in the tail and skin. Another limitation is that the experiments were carried out during a period of ∼2 years and at various hours during the day. Both factors might contribute noise to the thermal measurements. Nonetheless, we mitigated this variability using the ambient temperature compensation procedure.

## Conclusions

By using a multimodal approach for phenotyping social behavior in *Gtf2i^+/Del^* and *Gtf2i^+/Dup^* mice, we reveal a correlation between *Gtf2i* copy number variation and several behavioral and physiological variables in mice models of 7Dup and WBS. Thus, our work demonstrates, for the first time to our knowledge, the significant potential of thermography for phenotyping behavioral and physiological deficits in animal models of ASD. We believe that the phenotypes we found can be useful for future drug screening and for improved understanding of the phenotype associated with the *Gtf2i* gene.

## DECLARATIONS

### ETHICS APPROVAL

All experiments were approved by the University of Haifa Institutional Animal Care and Use Committee (IACUC) (Reference #: UoH-IL2301-103-4).

## CONSENT FOR PUBLICATION

Not applicable.

## DATA AVAILABILITY

The video dataset would be made available from the corresponding author upon reasonable request from an academic PI and will be free for academic use. DLC processed data, all statistical data (also found in supplementary file 1) for the figures, and processed data values used for the figures will be made available in a suitable public repository upon acceptance of the manuscript.

## AUTHOR CONTRIBUTIONS

D.P. – Data curation, Formal analysis, Investigation, Methodology, Writing – original draft, Software, Visualization.

S.N. – Data curation, Project administration, Resources, Software, Supervision.

N.R. - Data curation, Investigation.

T.S. – Data curation, Investigation.

S.W. Funding acquisition, Supervision, Writing – review and editing.

## Acknowledgements

ACKNOLEDGEMENTS

We thank Prof. Lucy Osborne and Prof. Giuseppe Testa for providing the *Gtf2i*^+/dup^ and *Gtf2i*^+/-^ mice. We thank Yaniv Goldstein, Janet Tabakova, Wjdan Awaisy, Shorook Amara and Ori Yacobi for their help annotating the videos and Sara Sheikh for drawing the experiment setup illustration.

## CODE AVAILABILITY

The code used for analysis will be made available in a suitable public repository upon acceptance of the manuscript.

