## Supplementary Figures for "Contrasting behavioral and physiological effects of *Gtf2i* duplication and deletion in mouse models of the 7q11.23 Duplication and Williams-Beuren Syndromes"

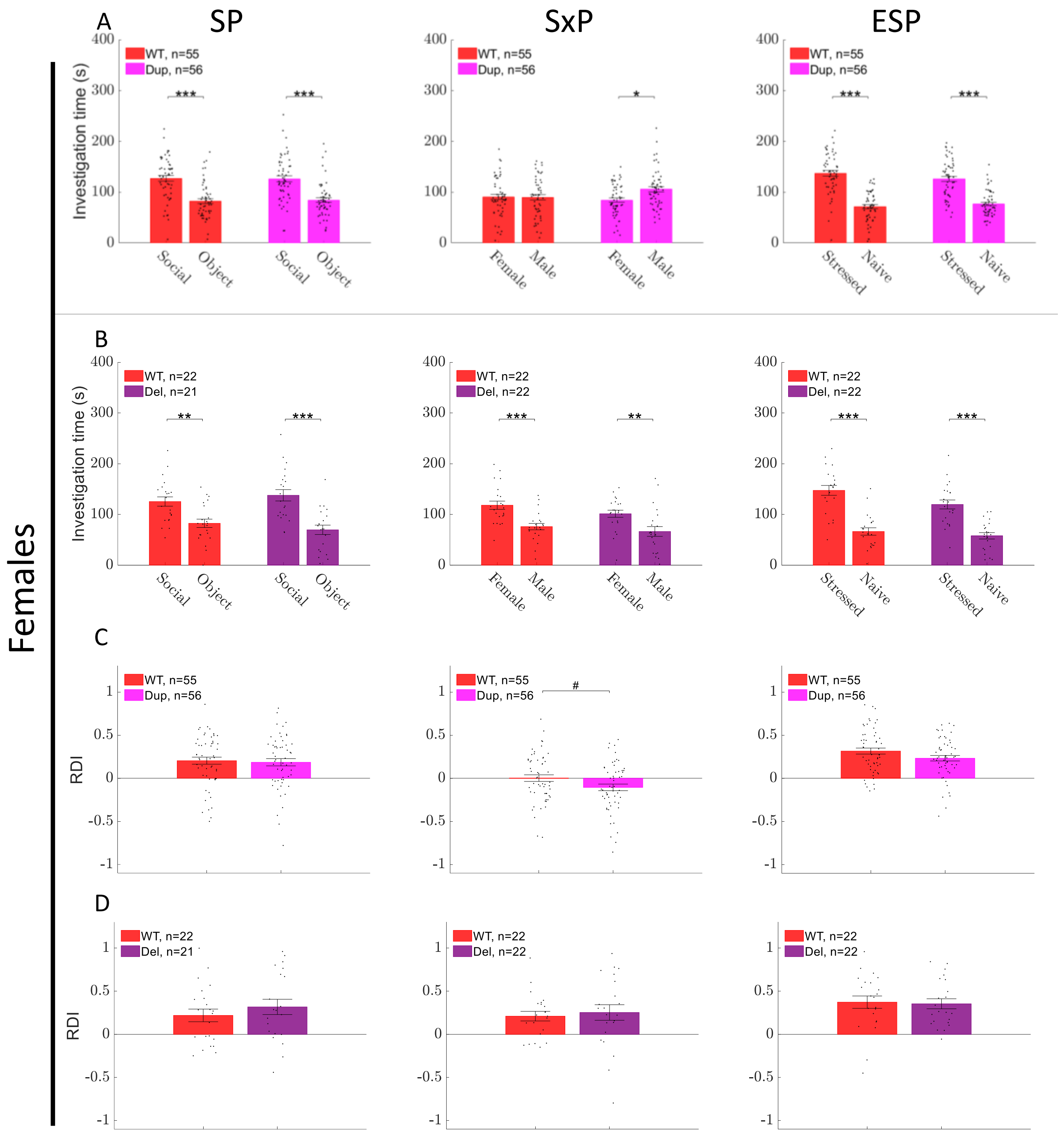


***Supplementary Fig. 1. Behavior in female mice.*** *Mean investigation time (±SEM) towards each of the two stimuli during SP, SxP, and ESP of* $WT_{Dup}$ *vs Dup (**A**),* $WT_{Del}$ *vs Del (**B**). (**C,D**) Shows the mean RDI (±SEM) towards stimulus1 (Social in SP, Female in SxP, and Stressed in ESP) for* $WT_{Dup}$ *vs Dup (**C**) and* $WT_{Del}$ *vs Del (**D**). Pairwise comparisons were done using a two-sided Wilcoxon rank sum test. (**A,B**) were FDR corrected (2 comparisons per panel). p-values equal to or smaller than 0.1,0.05,0.01, and 0.001 were marked by #, *, **, ***, respectively.*

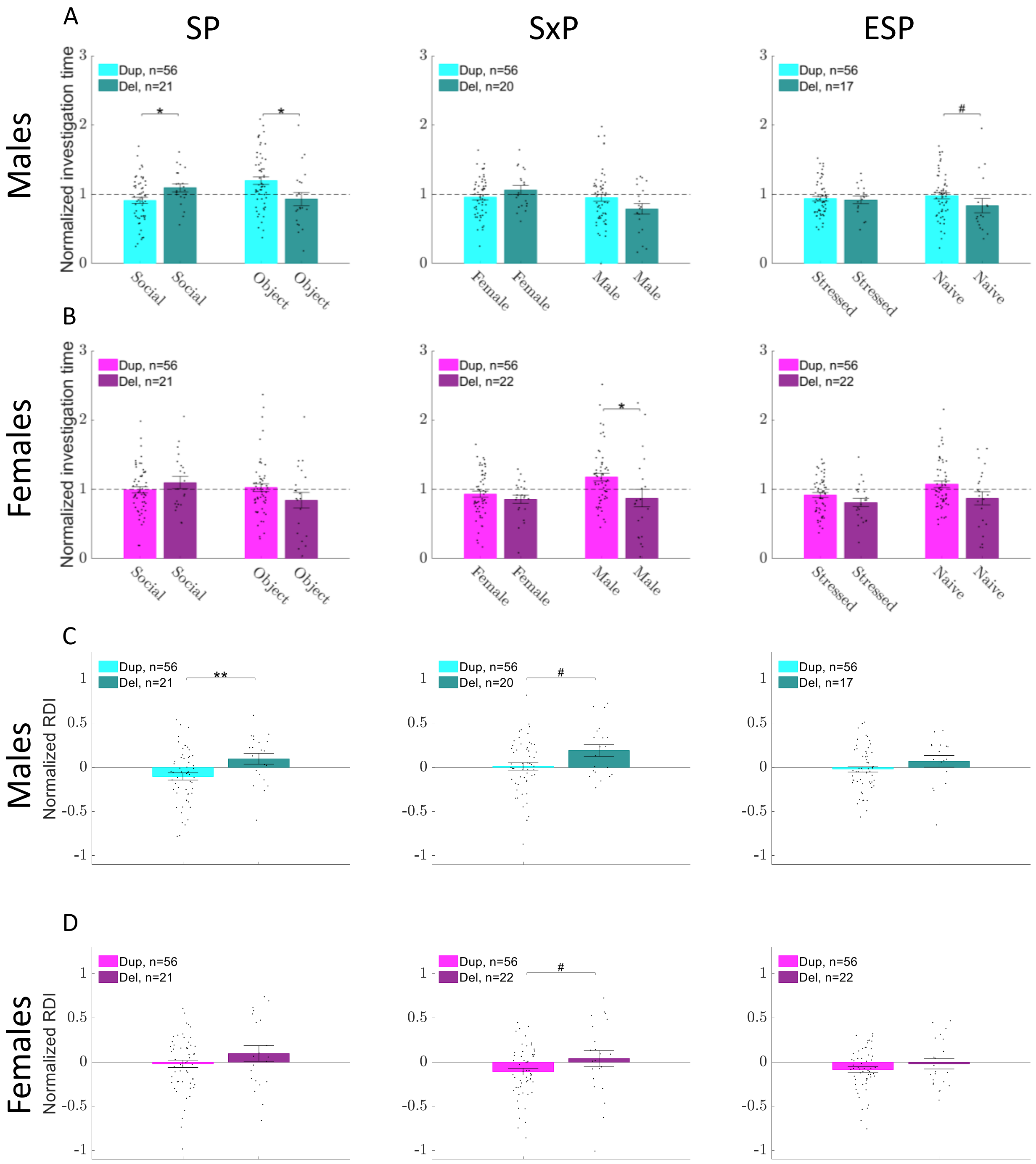


***Supplementary Fig. 2. Social discrimination in Dup vs Del.*** *(**A**) Shows the mean investigation time ±SEM towards each of the two stimuli during SP, SxP, and ESP in male Dup (Normalized by dividing by the mean of male $WT_{Dup}$) vs male Del (Normalized by dividing by the mean of male $WT_{Del}$). (**B**) Shows the same for females. (**C**) Shows comparisons between normalized RDI values of male Dup vs male Del. (**D**) Shows the same for females. Normalization of RDI values was done by subtracting the mean RDI value of $WT_{Dup}$ or $WT_{Del}$ from the RDI of Dup and Del, respectively. Pairwise comparisons were done using a two-sided Wilcoxon rank sum test. (**A,B**) were FDR corrected (2 comparisons per panel). p-values equal to or smaller than 0.1,0.05,0.01, and 0.001 were marked by #, *, **, ***, respectively.*

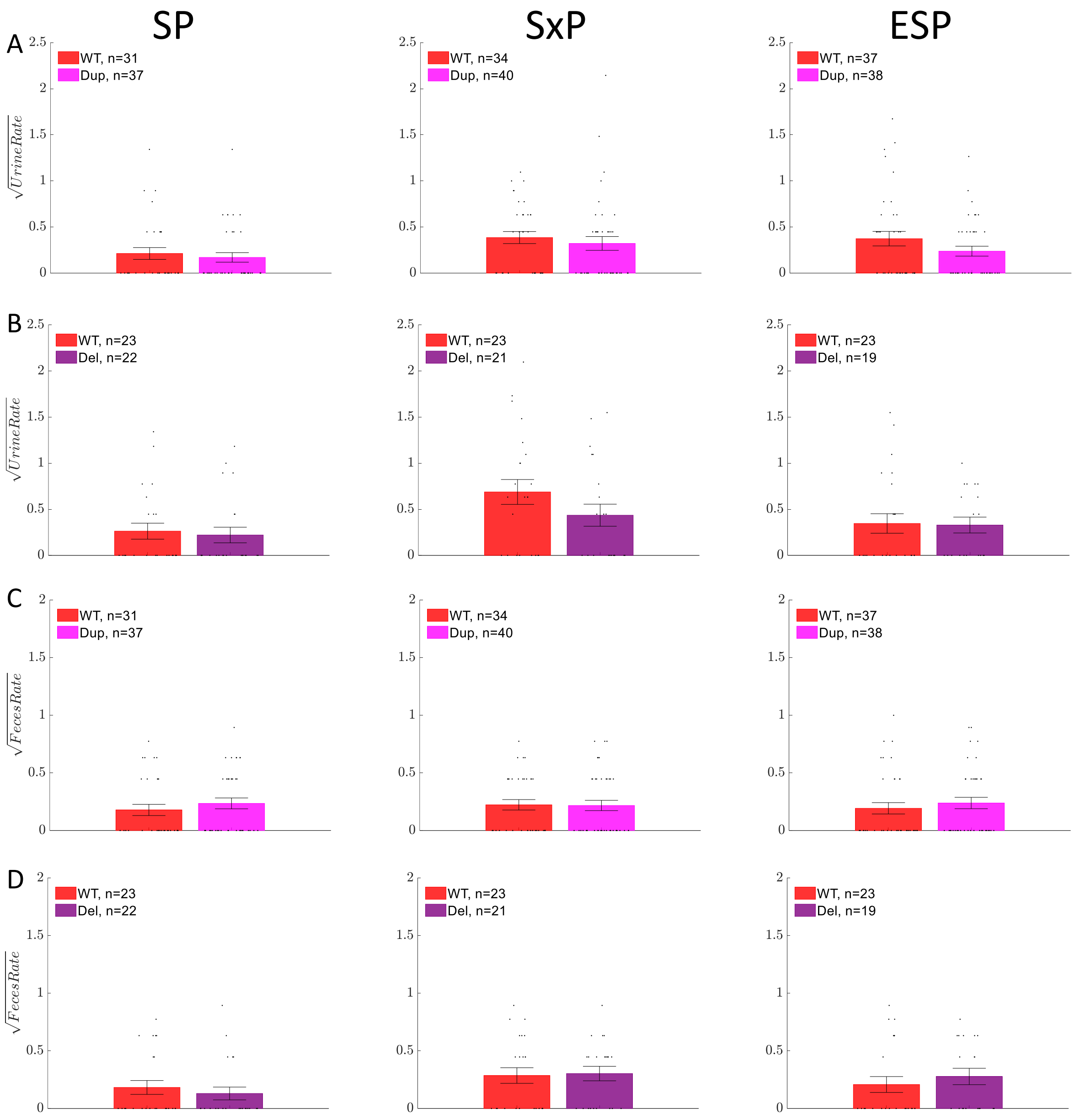


***Supplementary Fig. 3. Urination and defecation rates in female mice.*** *Square root of the # of urination spots per minute during SP, SxP, and ESP for* $WT_{Dup}$ *vs Dup (**A**),* $WT_{Del}$ *vs Del (**B**). (**C,D**) shows the same for defecation rate. Bars show the mean value, and error bars show ±SEM. For pairwise comparisons, we used two types of tests. A two-sided Wilcoxon rank-sum test with p-values equal to or smaller than 0.1,0.05,0.01, and 0.001 was marked by #, *, **, ***, respectively. A two-way Chi-square test (for comparing the number of zeros and non-zeros between two groups) with p-value smaller than 0.1,0.05,0.01, and 0.001 was marked by !,+,++,+++, respectively.*

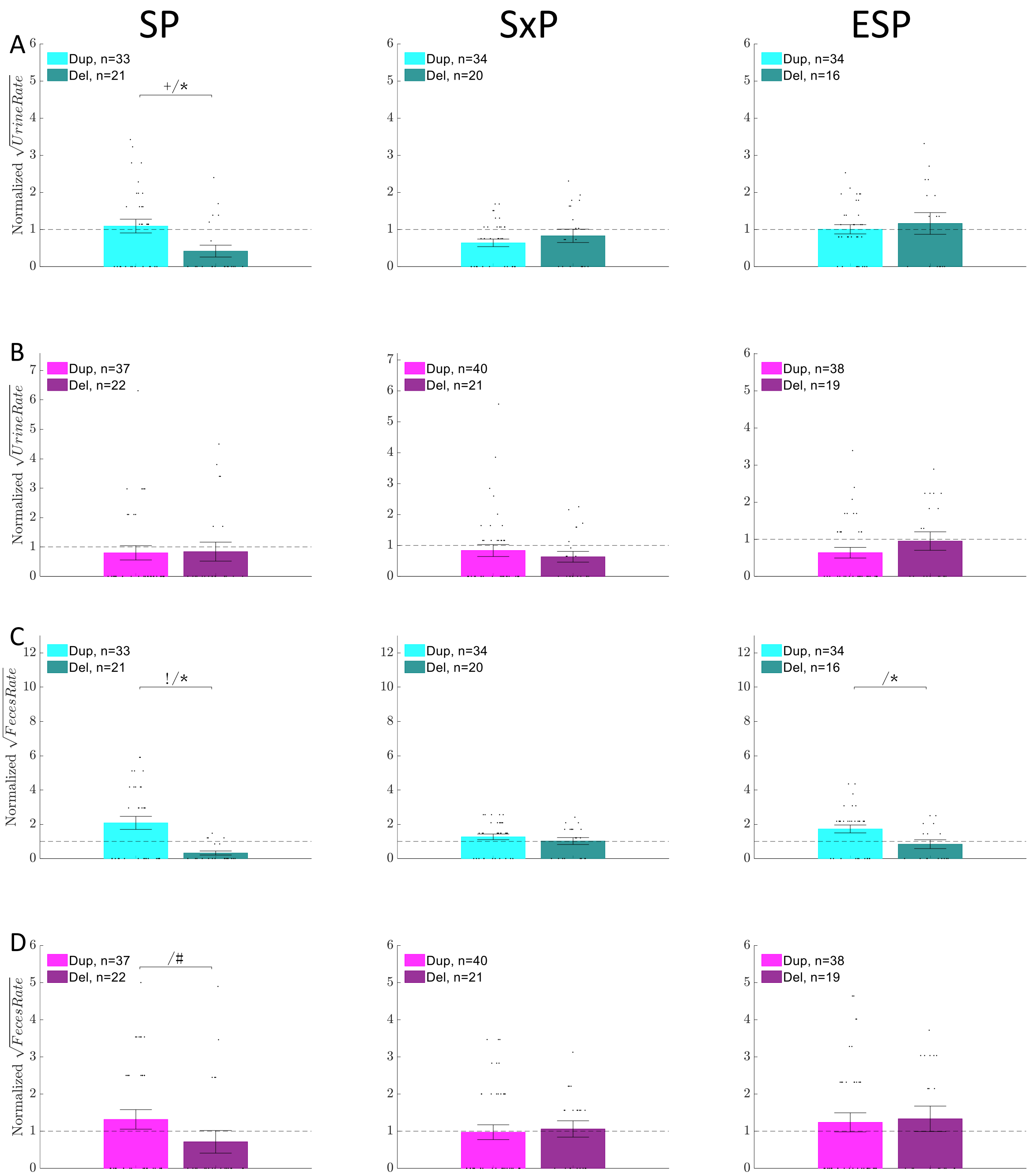


***Supplementary Fig 4. Urination and defecation rate in normalized Dup vs normalized Del.*** *Normalization was done by dividing Dup and Del values by the mean value of the relevant WT (square-rooted values were used). (**A, B**) compares the square root of urination rate (urine spots per minute) in Dup vs Del males (**A**) and Dup vs Del females (**B**). (**C, D**) shows the same for defecation rate. Bars show the mean value, and error bars show ±SEM. For pairwise comparisons, we used two types of tests. A two-sided Wilcoxon rank-sum test with p-values equal to or smaller than 0.1,0.05,0.01, and 0.001 was marked by #, *, **, ***, respectively. A two-way Chi-square test (for comparing the number of zeros and non-zeros between two groups) with p-value smaller than 0.1,0.05,0.01, and 0.001 was marked by !,+,++,+++, respectively.*

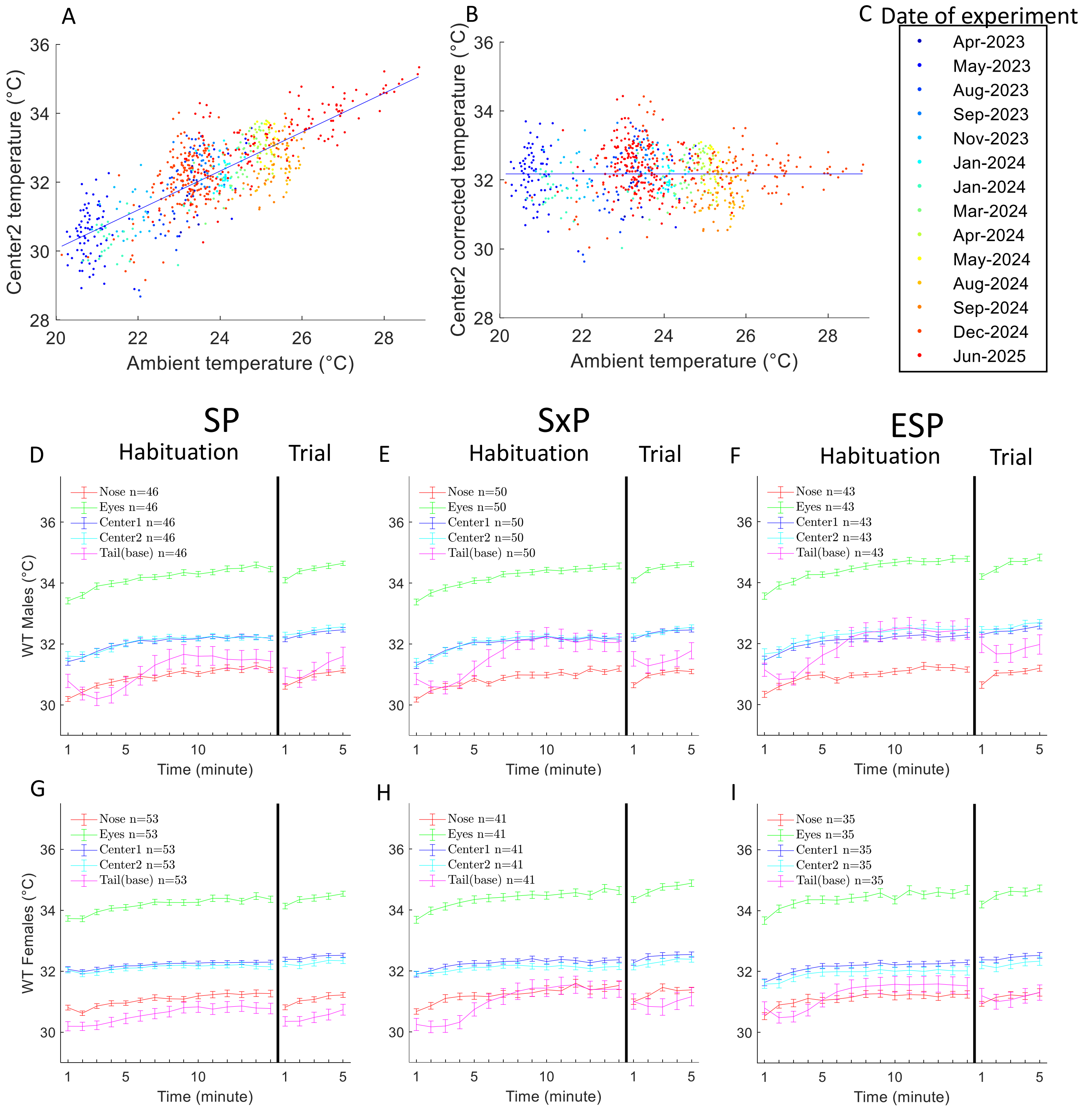


***Supplementary Fig. 5. Surface temperature measurements for mice body parts.*** *(**A,B,C**): Compensation function for the effect of the ambient temperature on the apparent temperature of the mice's body center (Center2 key-point). (**A**) Each dot marks the mean apparent temperature of a mouse at Center2 during habituation minutes 10-14 in a single experiment as a function of the ambient temperature in this experiment. Ambient temperature was estimated as the apparent temperature of the ambient temperature reference point (see Fig. 3D and Methods). The fitted blue line (linear regression) is used to compensate for the ambient temperature. Both male and female WT and mutant mice are included in the compensation function computation. This compensation reduces the Center2 measured temperature variance by 64.4% from 1.42°C to 0.51°C. Color shows the date at which the experiment took place according to the color legend in (**C**). (**B**) shows the values in (**A**) after compensation using the fitted line shown in (**A**). A separate compensation function was computed for each body part. (**D-I**): Mean temperature (±SEM) of several body parts in male mice (**D,E,F**) and female mice (**G,H,I**) throughout SP, SxP, and ESP tests for* $WT_{Dup}$ *and* $WT_{Del}$ *(pooled together).*

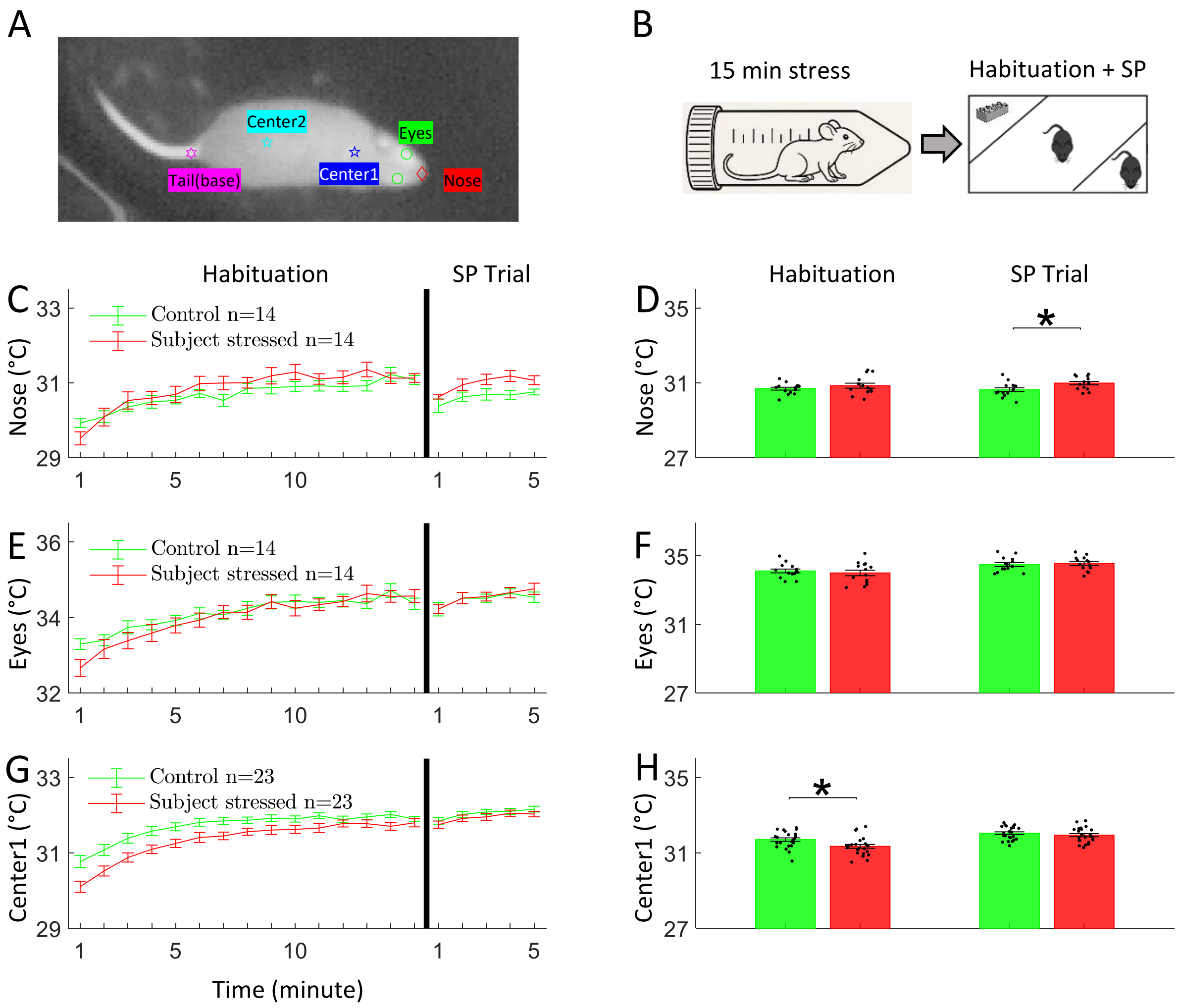


***Supplementary Fig. 6. Effect of subject stress on male mice temperature during SP test.*** *Location of the measure body parts location is shown in (**A**). Stress to the subject mouse is generated by holding it in a 50 ml plastic tube restrainer for 15 minutes (**B**) before the SP test. Mean temperature ±SEM along minutes 1-15 and trial minutes 1-5 for the Nose, Eyes and Center1 in* $WT_{Dup}$ *and* $WT_{Del}$ *pooled together is shown in (**C**,**E**,**G**) and quantized in (**D**,**F**,**H**). In each panel, the Control group and Subject stressed group includes the same mice. A two-sided Wilcoxon rank-sum test was used for pairwise comparisons. (**D**,**F**,**H**) were FDR corrected using Benjamini-Hochberg method (2 comparisons per panel). p-values equal to or smaller than 0.1,0.05,0.01, and 0.001 were marked by #, *, **, ***, respectively.*

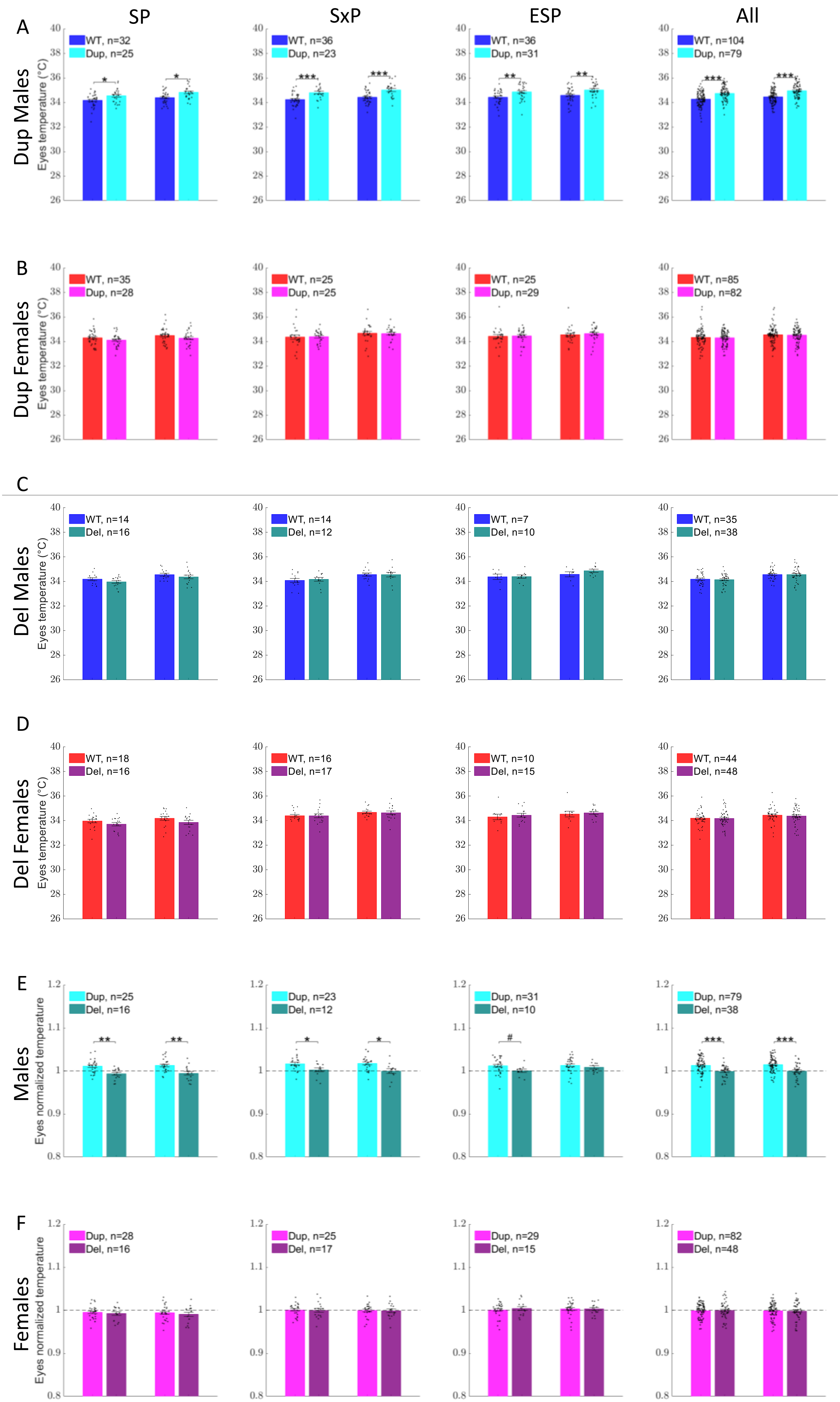

SP
SxP
ESP
All
A
Dup Males
Eyes temperature (°C)
WT, n=32
Dup, n=25
WT, n=36
Dup, n=23
WT, n=36
Dup, n=31
WT, n=104
Dup, n=79
B
Dup Females
Eyes temperature (°C)
WT, n=35
Dup, n=28
WT, n=25
Dup, n=25
WT, n=25
Dup, n=29
WT, n=85
Dup, n=82
C
Del Males
Eyes temperature (°C)
WT, n=14
Del, n=16
WT, n=14
Del, n=12
WT, n=7
Del, n=10
WT, n=35
Del, n=38
D
Del Females
Eyes temperature (°C)
WT, n=18
Del, n=16
WT, n=16
Del, n=17
WT, n=10
Del, n=15
WT, n=44
Del, n=48
E
Males
Eyes normalized temperature
Dup, n=25
Del, n=16
Dup, n=23
Del, n=12
Dup, n=31
Del, n=10
Dup, n=79
Del, n=38
F
Females
Eyes normalized temperature
Dup, n=28
Del, n=16
Dup, n=25
Del, n=17
Dup, n=29
Del, n=15
Dup, n=82
Del, n=48

***Supplementary Fig. 7. Eyes temperature across genotypes.*** *Eyes temperature during SP, SxP, ESP tests, and all tests pooled together for male* $WT_{Dup}$ *vs male Dup (**A**), female* $WT_{Dup}$ *vs female Dup (**B**), male* $WT_{Del}$ *vs male Del (**C**), female* $WT_{Del}$ *vs female Del (**D**), male Dup normalized to male* $WT_{Dup}$ *vs male Del normalized to male* $WT_{Del}$ *(**E**), female Dup normalized to female* $WT_{Dup}$ *vs female Del normalized to female* $WT_{Del}$ *(**F**). Bars show the mean temperature ±SEM. Pairwise comparisons were done using a two-sided Wilcoxon rank sum test, which were FDR corrected (2 comparisons per panel). p-values equal to or smaller than 0.1,0.05,0.01, and 0.001 were marked by #, *, **, ***, respectively.*

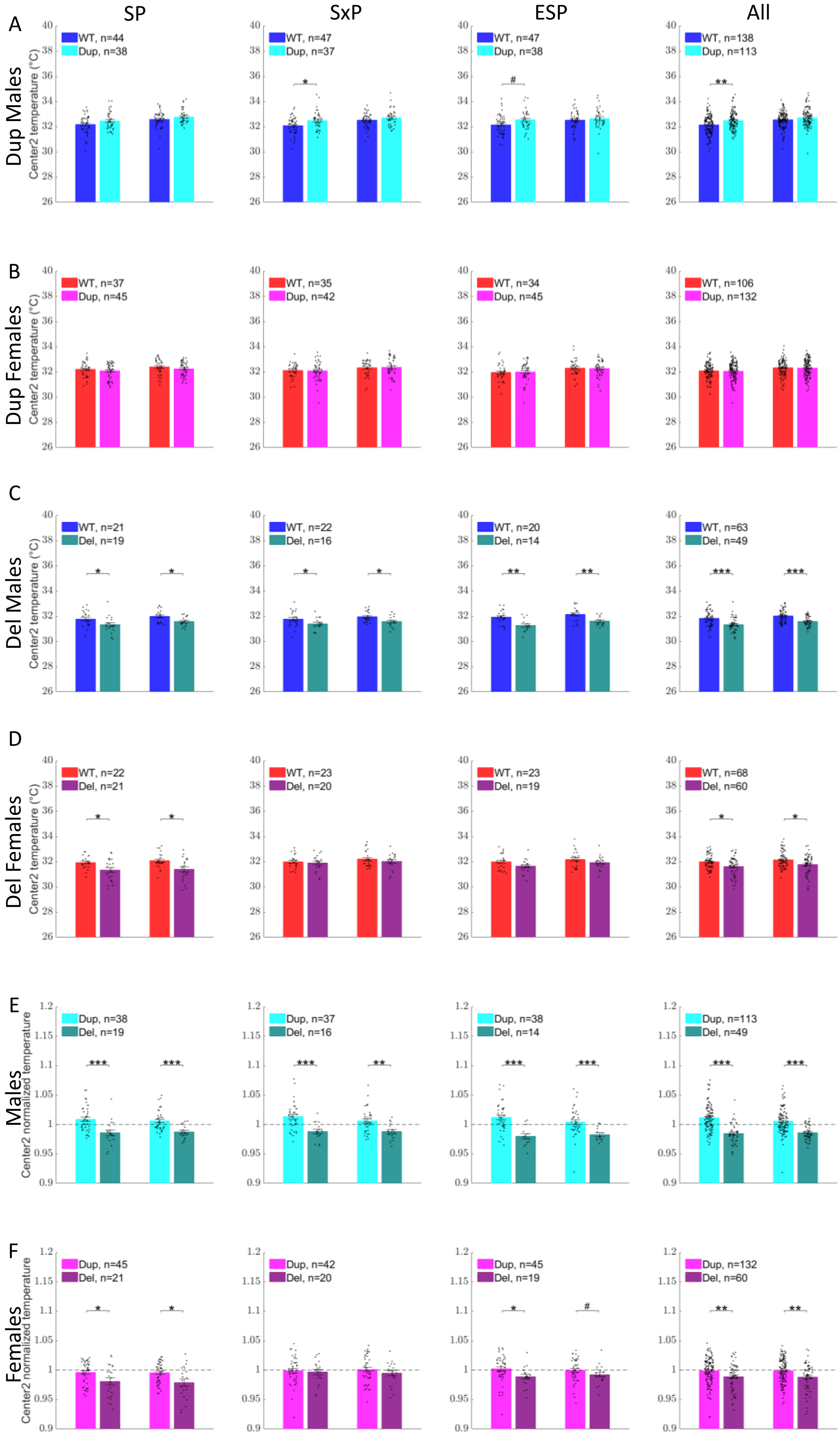
SP
SxP
ESP
All
A
Dup Males
Center2 temperature (°C)
WT, n=44
Dup, n=38
WT, n=47
Dup, n=37
WT, n=47
Dup, n=38
WT, n=138
Dup, n=113
B
Dup Females
WT, n=37
Dup, n=45
WT, n=35
Dup, n=42
WT, n=34
Dup, n=45
WT, n=106
Dup, n=132
C
Del Males
WT, n=21
Del, n=19
WT, n=22
Del, n=16
WT, n=20
Del, n=14
WT, n=63
Del, n=49
D
Del Females
WT, n=22
Del, n=21
WT, n=23
Del, n=20
WT, n=23
Del, n=19
WT, n=68
Del, n=60
E
Males
Center2 normalized temperature
Dup, n=38
Del, n=19
Dup, n=37
Del, n=16
Dup, n=38
Del, n=14
Dup, n=113
Del, n=49
F
Females
Dup, n=45
Del, n=21
Dup, n=42
Del, n=20
Dup, n=45
Del, n=19
Dup, n=132
Del, n=60

***Supplementary Fig. 8. Center2 temperature across genotypes.*** *Center2 temperature during SP, SxP, ESP tests, and all tests pooled together for male $WT_{Dup}$ vs male Dup (**A**), female $WT_{Dup}$ vs female Dup (**B**), male $WT_{Del}$ vs male Del (**C**), female $WT_{Del}$ vs female Del (**D**), male Dup normalized to male $WT_{Dup}$ vs male Del normalized to male $WT_{Del}$ (**E**), female Dup normalized to female $WT_{Dup}$ vs female Del normalized to female $WT_{Del}$ (**F**). Bars show the mean temperature ±SEM. Pairwise comparisons were done using a two-sided Wilcoxon rank sum test, which were FDR corrected (2 comparisons per panel). p-values equal to or smaller than 0.1,0.05,0.01, and 0.001 were marked by #, *, **, ***, respectively.*

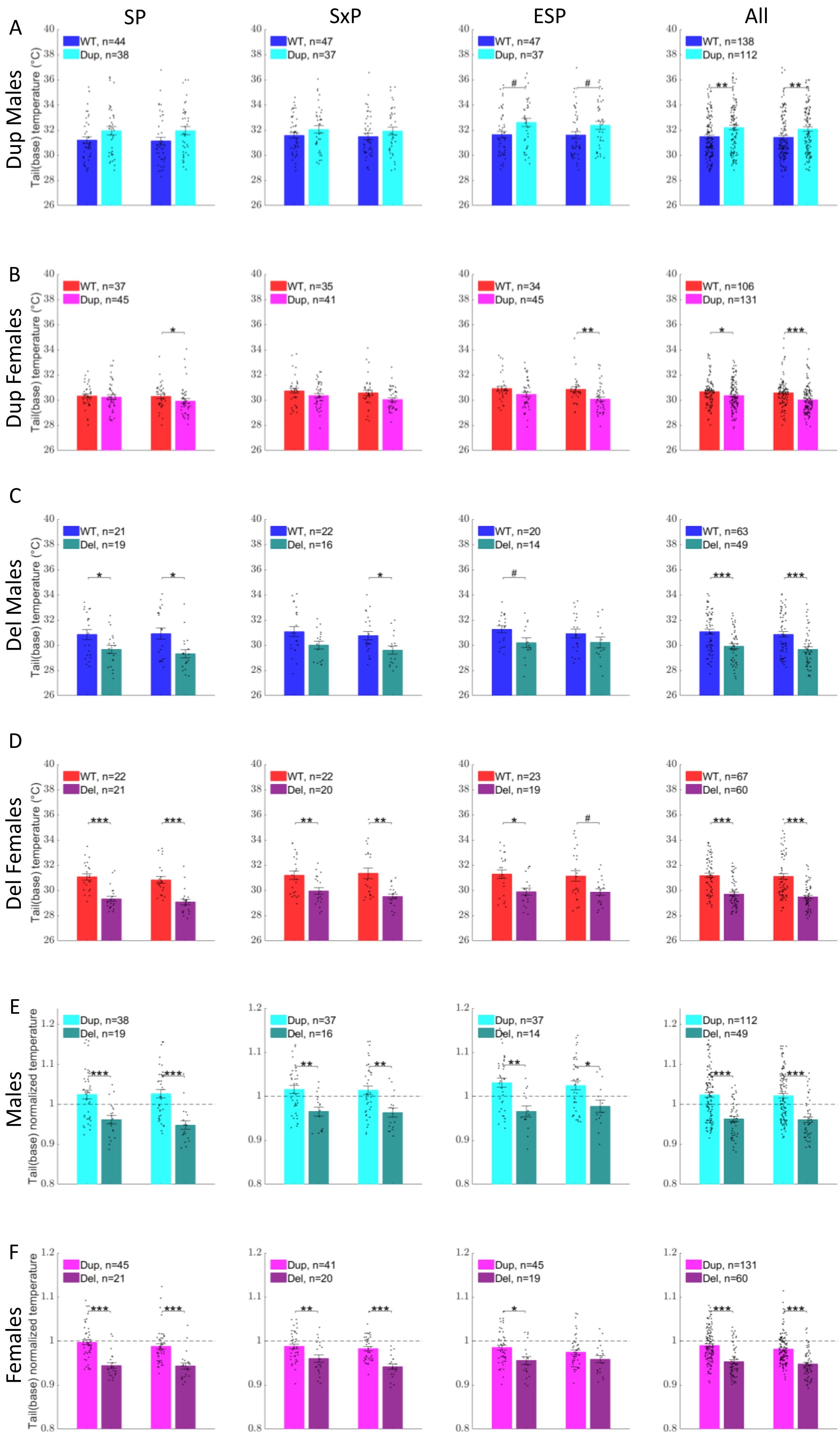

SP
SxP
ESP
All
A
Dup Males
B
Dup Females
C
Del Males
D
Del Females
E
Males
F
Females
Tail(base) temperature (°C)
Tail(base) normalized temperature
WT, n=44
Dup, n=38
WT, n=47
Dup, n=37
WT, n=47
Dup, n=37
WT, n=138
Dup, n=112
WT, n=37
Dup, n=45
WT, n=35
Dup, n=41
WT, n=34
Dup, n=45
WT, n=106
Dup, n=131
WT, n=21
Del, n=19
WT, n=22
Del, n=16
WT, n=20
Del, n=14
WT, n=63
Del, n=49
WT, n=22
Del, n=21
WT, n=22
Del, n=20
WT, n=23
Del, n=19
WT, n=67
Del, n=60
Dup, n=38
Del, n=19
Dup, n=37
Del, n=16
Dup, n=37
Del, n=14
Dup, n=112
Del, n=49
Dup, n=45
Del, n=21
Dup, n=41
Del, n=20
Dup, n=45
Del, n=19
Dup, n=131
Del, n=60

***Supplementary Fig. 9. Tail(base) temperature across genotypes.*** *Tail(base) temperature during SP, SxP, ESP tests, and All tests pooled together for male $WT_{Dup}$ vs male Dup (**A**), female $WT_{Dup}$ vs female Dup (**B**), male $WT_{Del}$ vs male Del (**C**), female $WT_{Del}$ vs female Del (**D**), male Dup normalized to male $WT_{Dup}$ vs male Del normalized to male $WT_{Del}$ (**E**), female Dup normalized to female $WT_{Dup}$ vs female Del normalized to female $WT_{Del}$ (**F**). Bars show the mean temperature ±SEM. Pairwise comparisons were done using a two-sided Wilcoxon rank sum test, which were FDR corrected (2 comparisons per panel). p-values equal to or smaller than 0.1,0.05,0.01, and 0.001 were marked by #, *, **, ***, respectively.*

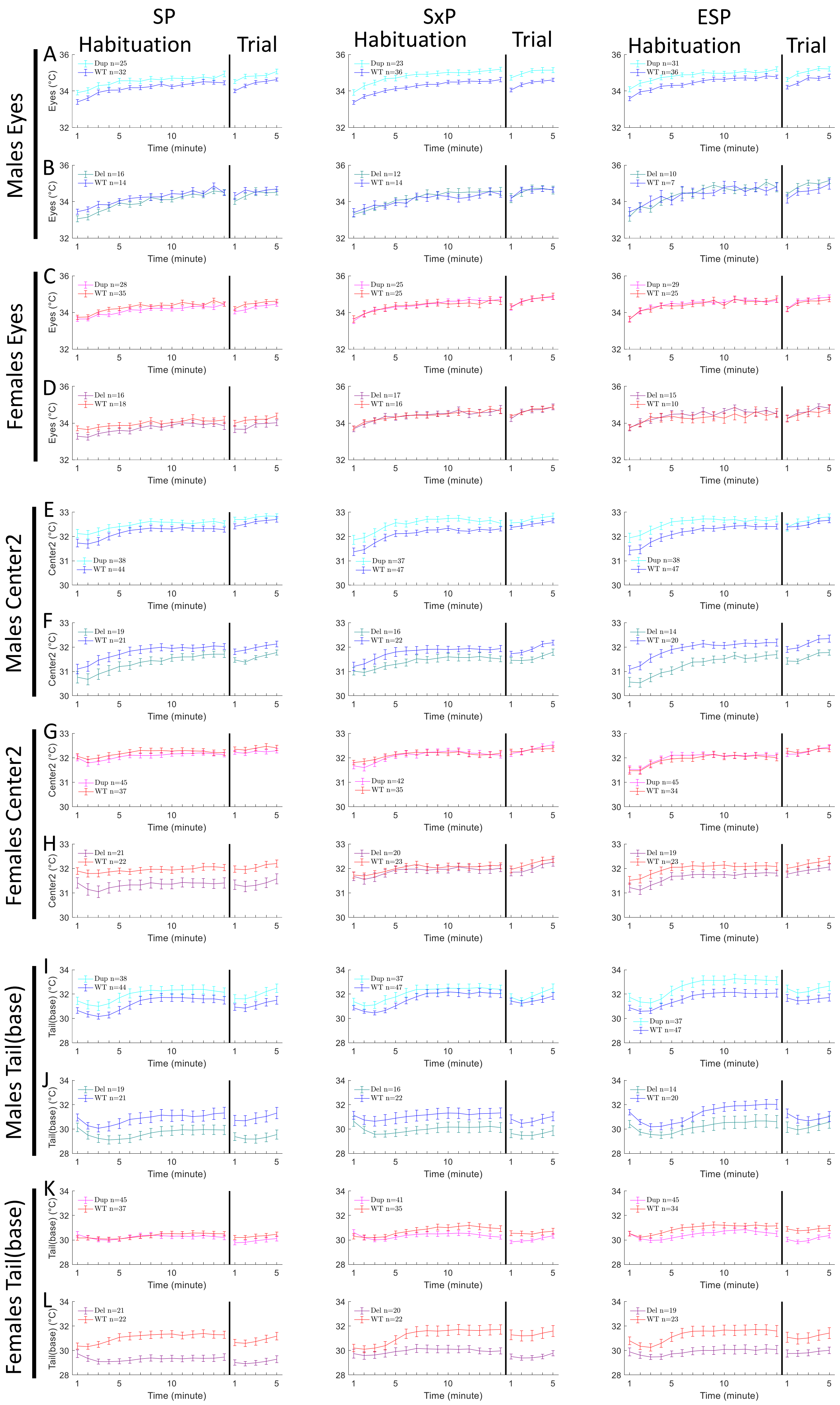
SP
SxP
ESP
Habituation
Trial
Males Eyes
Females Eyes
Males Center2
Females Center2
Males Tail(base)
Females Tail(base)
A
B
C
D
E
F
G
H
I
J
K
L
Eyes (°C)
Center2 (°C)
Tail(base) (°C)
Time (minute)
Dup n=25
WT n=32
Dup n=23
WT n=36
Dup n=31
WT n=36
Del n=16
WT n=14
Del n=12
WT n=14
Del n=10
WT n=7
Dup n=28
WT n=35
Dup n=25
WT n=25
Dup n=29
WT n=25
Del n=16
WT n=18
Del n=17
WT n=16
Del n=15
WT n=10
Dup n=38
WT n=44
Dup n=37
WT n=47
Dup n=38
WT n=47
Del n=19
WT n=21
Del n=16
WT n=22
Del n=14
WT n=20
Dup n=45
WT n=37
Dup n=42
WT n=35
Dup n=45
WT n=34
Del n=21
WT n=22
Del n=20
WT n=23
Del n=19
WT n=23
Dup n=38
WT n=44
Dup n=37
WT n=47
Dup n=37
WT n=47
Del n=19
WT n=21
Del n=16
WT n=22
Del n=14
WT n=20
Dup n=45
WT n=37
Dup n=41
WT n=35
Dup n=45
WT n=34
Del n=21
WT n=22
Del n=20
WT n=22
Del n=19
WT n=23

***Supplementary Fig. 10. Gtf2i dosage effect on mice surface temperature.*** *Eyes temperature for Dup and* $WT_{Dup}$ *males (**A**), Del and* $WT_{Del}$ *males (**B**), Dup and* $WT_{Dup}$ *females (**C**), and Del and* $WT_{Del}$ *Females (**D**) throughout SP, SxP, and ESP tests. (**E-H**) shows the same for Center2. (**I-L**) shows the same for Tail(base). The plots show the mean temperature ±SEM per minute during minutes 1-15 of the habituation and minutes 1-5 of the trial.*

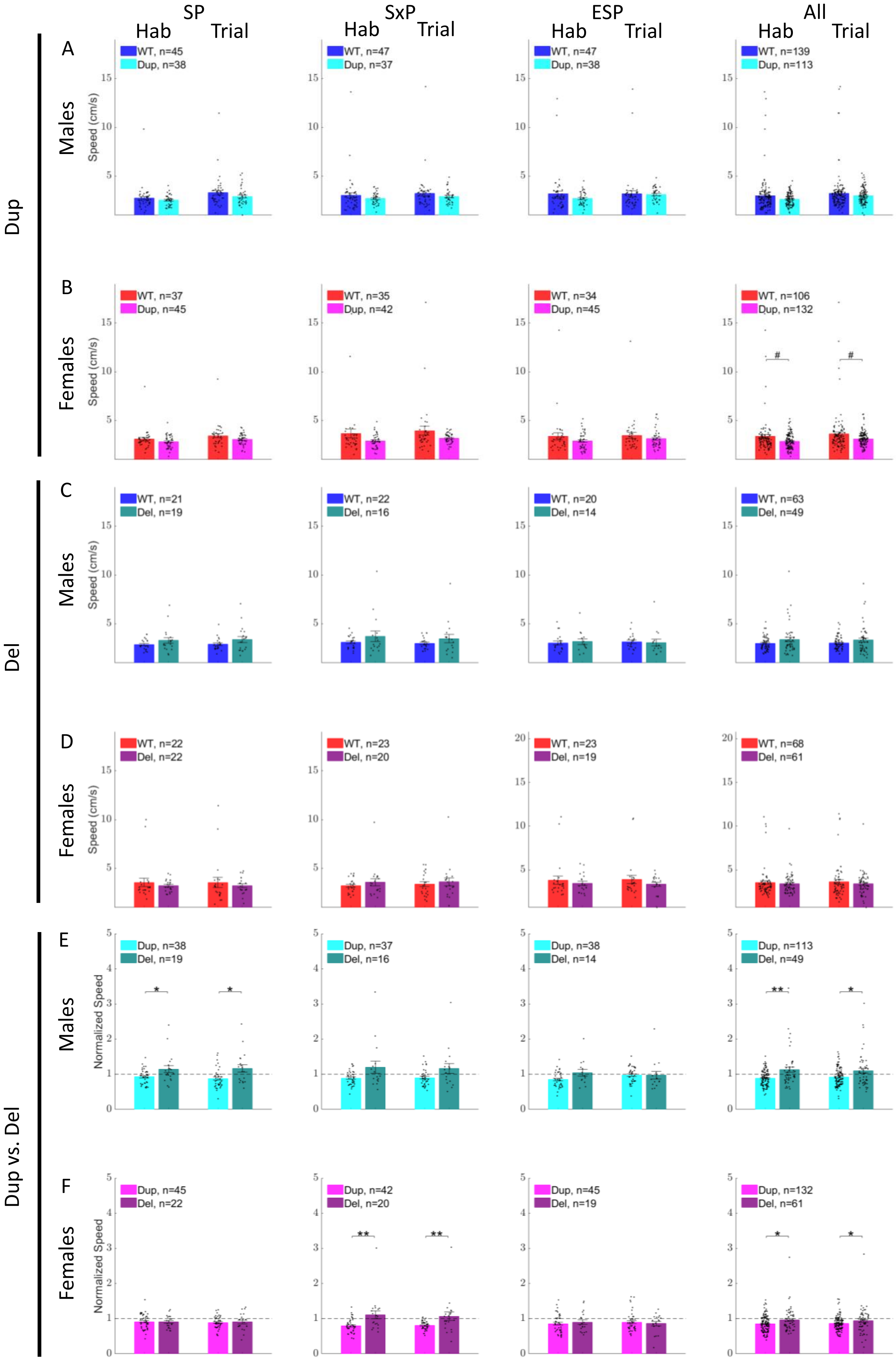

SP
SxP
ESP
All
Hab
Trial
Dup
Del
Dup vs. Del
A
B
C
D
E
F
Males
Females
Speed (cm/s)
Normalized Speed
WT, n=45
Dup, n=38
WT, n=47
Dup, n=37
WT, n=47
Dup, n=38
WT, n=139
Dup, n=113
WT, n=37
Dup, n=45
WT, n=35
Dup, n=42
WT, n=34
Dup, n=45
WT, n=106
Dup, n=132
WT, n=21
Del, n=19
WT, n=22
Del, n=16
WT, n=20
Del, n=14
WT, n=63
Del, n=49
WT, n=22
Del, n=22
WT, n=23
Del, n=20
WT, n=23
Del, n=19
WT, n=68
Del, n=61
Dup, n=38
Del, n=19
Dup, n=37
Del, n=16
Dup, n=38
Del, n=14
Dup, n=113
Del, n=49
Dup, n=45
Del, n=22
Dup, n=42
Del, n=20
Dup, n=45
Del, n=19
Dup, n=132
Del, n=61

***Supplementary Fig. 11. Movement speed during tests.*** *Mean speed (cm/second) ±SEM during habituation (minutes 1-15 of habituation) and during the trial periods (minutes 1-5 of the trial) are shown for male $WT_{Dup}$ vs male Dup (**A**), female $WT_{Dup}$ vs female Dup (**B**), male $WT_{Del}$ vs male Del (**C**), female $WT_{Del}$ vs female Del (**D**), male Dup normalized to male $WT_{Dup}$ vs male Del normalized to male $WT_{Del}$ (**E**), female Dup normalized to female $WT_{Dup}$ vs female Del normalized to female $WT_{Del}$ (**F**). The "All" column pools the SP, SxP, and ESP tests together. Pairwise comparisons were done using a two-sided Wilcoxon rank sum test, which were FDR corrected (2 comparisons per panel). p-values equal to or smaller than 0.1,0.05,0.01, and 0.001 were marked by #, *, **, ***, respectively.*
