## Supplementary File 1 for "Contrasting behavioral and physiological effects of *Gtf2i* duplication and deletion in mouse models of the 7q11.23 Duplication and Williams-Beuren Syndromes": Readme.docx

### Supplementary Data and Statistical Information

#### Video database:

The list of thermal videos included in this paper is listed in SupplementaryTable1-VideoDatabase.csv. The first column of the CSV file contains the video folder and video name. The video name contains the following information:

<cage ID and gender(M/F)>_<mouse index>_<Test Type: SP or SxP or Stress(ESP)>_<ICR Dup or ICR Del genotype background>_<WT or HET(mutant)>_<time and date>

The ICR Dup is mentioned in the video name for both Dup mice and WT_Dup_ mice.

The ICR Del is mentioned in the video name for both Del mice and WT_Del_ mice.

The cage ID is a unique identifier of the cage within the group of ICR Dup and within the group of ICR Del videos, but it is possible that an ICR Dup cage and an ICR Del cage will have the same cage ID.

The rest of the columns include the following information

*vidType*: either SP or SxP or Stress (ESP)

*isMale*: 1 for male and 0 for female

*isWT*: 1 for WT and 0 for mutant (either Dup or Del)

*isDup*: 1 for Dup mutation (3 copies of Gtf2i), 0 otherwise

*isDel*: 1 for Del mutation (1 copy of Gtf2i), 0 otherwise

*isWT_Dup*: 1 for WT_Dup_, 0 otherwise

*isWT_Del*: 1 for WT_Del_, 0 otherwise

*ValidForUrineFeces:* 1 if this video was used in the urination and defecation rate figures. 0 otherwise.

*ValidForThermal:* 1 if this video was used in the thermal measurement and mean speed figure. 0 otherwise.

*remarks:* remarks regarding this video. Usually contains the reason for excluding this video.

#### Data and statistics:

The data points and statistical information for all of the figures that include statistical comparisons are included in Excel files in the following sub-directories:

Fig1_And_SupFig1-2_BehaviorAndRDI

Fig2_And_SupFig3-4_UrineFeces

Fig3-5_And_SupFig5-9_Temperature

SupFig11_MeanSpeed

Each Excel file includes several sheets. The sheet name includes the gender and the test type: SP, SxP, ESP, and All (All means SP, SxP, and ESP pooled together).

Each sheet contains the following information for each group of values:

1. Mean, median, std, sample size, and the list of raw values.
2. Statistical information for each comparison:
   a. *pVal (ranksum) and Statistical_Significance_Symbol* (#,*,**,***) which matches p-values smaller than or equal to 0.1, 0.05, 0.01, 0.001.
   b. *z_statistics and ranksum_statistics*: statistics related to the pVal computation (generated by Matlab’s ranksum function, see Matlab’s documentation).
   c. *FDR*: p-value corrected by FDR (if applicable).

##### Additional remarks:

**Urination and Defecation data in Fig2_And_SupFig3-4_UrineFeces sub-directory**

The Urination and Defecation Excel files contain additional information regarding the comparison of the distribution of zeros in the urination and defecation data.

*ZeroDistributionPval*: p-value of the two-way chi-square test comparing the number of zeros in each group. Significance symbols are !,+,++,+++ for p-value smaller than 0.1,0.05,0.01,0.001

*ZeroDistributionChiStat*: chi-square test statistic

The urination and defecation Excel files include a separate Excel file in which the raw values have been square-rooted (as was used in Fig. 2. and Supplementary Fig. 3) and an Excel file that contains the original raw values without the square root. Please note “WithSqrt” or “NoSqrt” text in the folder name.
